# Targeting the Human Papillomavirus 16 E6 Oncoprotein with Antibodies

**DOI:** 10.1101/2022.10.21.513107

**Authors:** Guillem Dayer, Ashley Faulkner, Tanu Talwar, Melissa Togtema, Ingeborg Zehbe

## Abstract

The human papillomavirus (HPV) 16 genome encodes two oncoproteins, E6 and E7, which are essential for viral carcinogenesis. While E7 promotes cell proliferation, E6 abolishes the resulting p53-dependent apoptotic response. Due to this specific function, E6 is considered a suitable target for the development of a variety of therapeutic agents such as antibodies. Here, we review anti-E6 antibodies/antibody fragments generated by us and others, as well as present our latest results with *Camelidae*-derived single-domain antibodies (sdAbs). We had previously isolated a pool of anti-E6 sdAbs to identify E6 binders with the potential to be used clinically and in research. While our previous work has focused on recombinant E6 proteins, here we evaluated these sdAbs’ binding capacity to the endogenous E6 protein using co-immunoprecipitation and immunofluorescence. We obtained reproducible results in these applications with two sdAbs, filling a gap in HPV research. Despite their apparent E6 binding ability, these sdAbs do not raise p53 levels or induce apoptosis. Thus, while these reagents are valuable diagnostic and detection tools, identifying their therapeutic potential will require further development and testing.

## Introduction

The differences in the pathophysiology of the diseases caused by low- and high-risk human papillomaviruses (HPVs) are directly linked to functional differences in their E6 and E7 oncoproteins. These mainly include the E6- or E7-dependent degradation of host cellular proteins leading to viral infection, persistence, and tumorigenesis (Ghittoni et al. 2010). While E7 triggers aberrant cell proliferation by degrading pRB, the E6 protein inactivates the resulting p53-dependent apoptotic response by degrading the p53 protein in an E6AP-dependent manner. HPV infection can be prevented through vaccination, but reliable tools to detect or treat it are still inadequately addressed. HPV16 is the type, most commonly found in cervical carcinomas. Due to its anti-apoptotic activity, HPV16 E6 has been a candidate for therapeutical development (Hoppe-Seyler et al. 2018 and references therein). It has been reasoned that targeting E6 will restore p53 activity leading to apoptosis in HPV-infected but not in normal cells (Togtema et al. 2012, Togtema et al. 2019). Several molecules aiming at abolishing E6 functions have been generated, such as chemical compounds (*i*.*e*., zinc-ejector compounds), small interfering ribonucleic acids (siRNAs), RNA aptamers, antibodies (Abs), and inhibitory peptides (reviewed in Togtema et al. 2019). Abs against HPV16 E6 such as full-sized monoclonal Abs (mAb), single-chain variable fragment Abs (scFvs) and single-domain Abs (sdAbs) were primarily designed for therapeutic purposes (Giovane et al. 1999, Masson et al. 2003, Lagrange et al. 2005, 2007, Griffin et al. 2006, Amici et al. 2016, Togtema et al. 2019, Zhang et al. 2021) but no molecule has yet reached clinical trials. The imaging of HPV16 detection by Abs in intact cells may be useful for research studies or as prognostic factors in the clinic, but these approaches are also currently limited by the lack of adequate reagents.

Our lab has recently isolated a set of llama-derived sdAbs (also referred to as VHHs or nanobodies) whose binding ability to HPV16 E6 was tested with recombinant E6 proteins (Togtema et al. 2019). Here, we show data that these sdAbs target endogenous E6 proteins in cancer-derived cell lines containing HPV16 genomic material, using lysed or intact cells. We begin with a literature overview of Abs generated against HPV16 E6, exploring their selection process (summarized in Table S1) and evaluation (summarized in Table S2) with emphasis on sdAbs. Secondly, we provide experimental evidence for the functionality of our sdAbs in immunodetecting endogenous E6: we identified two sdAbs capable of binding to E6 but without the desired effect of raising p53 levels or inducing apoptosis. These new reagents may be useful for HPV16 E6 detection but require further development and testing for therapeutic purposes.

## Materials and Methods

### Recombinant proteins

Three recombinant E6 proteins derived from HPV16 E6 as well as the maltose-binding protein (MBP) were used. For consistency, we adopted the earlier reported recombinant protein nomenclatures (see below). The tag-free recombinant E6 containing the amino acid substitution C23S, C58S, C87S, C104S, C118S, C147S was obtained from BostonBiochem (Cat. # AP-120). This recombinant protein is the commercially available version of the E6 6C/6S previously described (Nominé et al. 2001, Zanier et al. 2010). The tagged recombinant E6 proteins were expressed from plasmids kindly provided to us by Dr. Gilles Travé. These E6 proteins were engineered to include an hexahistidine (His6) tag, an MBP tag, and 4 cysteine/serine mutations to enhance their solubility. The tags at the N-terminal end of the protein are referred to as: His6MBP-4C/4S E6 and His6MBP-F47R 4C/4S E6. The His6MBP-4C/4S E6 contains mutations C80S, C97S, C111S, C140S and His6MBP-F47R 4C/4S E6 contains F47R, C80S, C97S, C111S, C140S (Togtema et al. 2019). MBP was purchased from Abnova (Cat. # P4989).

### Cell culture

Mammalian cell cultures, CaSki (ATCC CRL-1550), SiHa (ATCC HTB-35) (both HPV16-positive) and C33A (ATCC HTB-31) (HPV16-negative) were cultured in T75 flasks (Fisher Scientific, Cat. # 1368065) and maintained in a 5% CO_2_, 37°C humidified incubator in Dulbecco’s Modified Eagle Medium (DMEM, Fisher Scientific, Cat. # SH3024301) supplemented with 10% heat-inactivated fetal bovine serum (FBS, Fisher Scientific, Cat. # SH3039603) and 1% antibiotic/antimycotic (Fisher Scientific, Cat. # SV3007901). Cells were passaged to maintain 70-80% confluency and routine screening for *Mycoplasma* contamination was regularly performed. For subculture, cells were rinsed with 10 mL DPBS (DPBS/MODIFIED, Fisher Scientific, Cat. # SH30028.02) before adding 3 mL trypsin (Fisher Scientific, Cat. # SH3023602) for 10 minutes. Trypsin was neutralized with 3 volumes of complete DMEM and centrifuged at 125 x g for 5 minutes to pellet the cells. For cell culture, 2-5 × 10^5^ cells were reseeded. For cell harvesting, cells were rinsed with DPBS and centrifuged at 350 x g for 5 minutes twice, DPBS was removed, and the cell pellets were stored at -80°C.

### Dot blot

For recombinant E6 proteins and sdAbs: 2 µg of E6 (R&D Systems Inc., Cat. # AP-120-025), MBP (Abnova, Cat. # P4989), His6MBP-F47R 4C/4S E6 and His6MBP-4C/4S E6 were spotted on a 0.45 µm nitrocellulose membrane (BioRad, Cat. # 162-0116) and allowed to dry at room temperature for 1 hour. Once dry, each membrane was incubated in blocking solution for 1 hour at room temperature on a tube rotator. The blocking solution consisted of 10 mL TBST 0.01% and 0.5 g instant skim milk powder (MTBST 0.01%). Following blocking, membranes were cut and divided into respective 50 mL Falcon tubes containing primary antibody solution. The primary antibody solutions consisted of 5 mL TBST 0.01% and 5.4 μg/mL of the respective sdAb as previously described (Togtema et al. 2019). Membranes in primary antibody solution were incubated overnight at 4°C on a tube rotator. After incubation, membranes were washed 3 x with 5 mL of TBST 0.01%. Following washes, membranes were incubated in secondary antibody solution overnight at 4°C on a tube rotator. The secondary antibody solution consisted of MonoRab ™: Rabbit Anti-Camelid VHH Antibody [HRP] mAb (GenScript, Cat. # A01861-200) (1:5000) in MTBST 0.01%. Following incubation, membranes were washed 3 x with 5 mL of TBST 0.01%. Once washed, chemiluminescent solution (Clarity™ Western ECL Substrate, BioRad, Cat. # 170-5061) was applied and membranes were visualized using a UVP™ Gel Imaging System and VisionWorks software.

Dot blots were also performed to confirm the binding of sdAbs to Affi-Gel resin (as described in the *Co-immunoprecipitation* section) in co-immunoprecipitation experiments. Here, 2 μL of input and unbound sample were spotted on nitrocellulose membrane and allowed to dry at room temperature for 30 minutes. Once dry, the membrane was incubated in blocking buffer (MTBST 0.05%) for 30 minutes at room temperature on a tube rotator. The membrane was then incubated in antibody detection solution consisting of 1:20000 anti-HA rat mAb conjugated to horseradish peroxidase (HRP) (Roche [3F10], Cat. # 1201381900) diluted in MTBST 0.05% for 1 hour at room temperature on a tube rotator. The membrane was then washed 3 × 5 min with 5mL TBST 0.05% at room temperature on a tube rotator and imaged as described above.

### Cell lysis

Cell pellets were frozen at -80°C for at least 1 hour prior to lysis. Lysis buffer composed of Mammalian Protein Extraction Reagent (Fisher Scientific, Cat. # PI78501), 150 mM sodium chloride (NaCl, Fisher Scientific, Cat. # S271-3), 1X HALT™ phosphatase inhibitor cocktail (Fisher Scientific, Cat. # PI78428), 1X cOmplete EDTA free protease inhibitor (Roche, Cat. # 04693159001), and 1 mM phenylmethylsulphonyl fluoride was mixed and chilled at 4°C. Each fresh pellet was resuspended in 10 µL per mg of pellet weight of chilled lysis buffer and incubated on a tube rotator for 30 minutes at 4°C. The solutions were centrifuged at 14000 x g for 15 minutes at 4°C to remove cell debris and obtain the protein containing supernatant. A Bradford assay was performed to determine protein concentration.

### Western blot

Samples were resuspended in 1X SDS loading buffer with reducing agent (2 mM dithiothreitol (DTT), Fisher Scientific, Cat. # R0861) and heated to 95°C for 10 minutes before being loaded into precast gels (Mini-PROTEAN® TGX™ Precast Protein Gels, BioRad, Cat. # 4561094, 4561096). The Precision Plus Protein™ Dual Color Standards (BioRad, Cat. # 1610374) was used as the protein ladder. Gels were run in 4°C running buffer (14.4 g glycine, 3.03 g Tris Base, and 0.5 g SDS) at 120V for 75 minutes. Following the completion of the run, the proteins were transferred to a methanol activated PVDF membrane (Fisher Scientific, Cat. # PI88518). Electroblotting was done at 100V, 0.35A for 1 hour in chilled transfer buffer (14.4 g glycine, 3.03 g Tris Base and 20% methanol). Following the transfer, membranes were blocked with MTBST 0.05% for 1 hour at room temperature on a tube rotator. Following blocking, membranes were incubated with primary antibody overnight at 4°C on a tube rotator. E6 detection was performed using the 6F4 mAb, a kind a gift from Arbor Vita Corporation (Fremont, CA, USA), using a concentration of 2.69 µg/mL in MTBST 0.05%. For recombinant protein (His6MBP-4C/4S E6, His6MBP-F47R 4C/4S E6, E6 6C/6S and MBP) detection, 2.69 µg/mL of C26 or 6F4 in MTBST 0.05% was used as previously described (Togtema et al. 2019). Following overnight incubation, membranes were washed 3 times for 5 minutes in TBST 0.05% before incubation with the secondary antibody diluted in MTBST 0.05% for 1 hour at room temperature on a tube rotator. 6F4 was probed using a goat anti-mouse antibody conjugated to HRP 1:1000 (Jackson ImmunoResearch Laboratories Inc., Cat. # 115-035-062). SdAb detection was performed using (1:20000) MonoRab™ Rabbit Anti-Camelid VHH Antibody conjugated to HRP (GenScript, Cat. # A01861-200) in MTBST 0.05%. Membranes were washed for 3 × 5 minutes on a tube rotator with 5mL of TBST 0.05%. Once washed, chemiluminescent solution (Clarity Western ECL Substrate, Bio-Rad, Cat. # 1705061) was applied, and membranes were visualized using a UVP™ Gel Imaging System and VisionWorks software.

### Co-immunoprecipitation

Affi-Gel resin (BioRad, Cat. # 1536099 (Affi-Gel 10), Cat. # 1536051 (Affi-Gel-15)) was aliquoted (12.5 µL pure resin per column) and resuspended using HPLC-grade distilled water (Fisher Scientific, Cat. # W5-4). Affi-Gel resin was divided equally into Pierce™ Spin Columns (Fisher Scientific, Cat. # PI69725). Unless otherwise specified, all centrifugations were performed for 30 seconds at 1000 x g at 4°C and all incubations were performed at 4°C on a tube rotator. Columns were centrifuged and the resin storage buffer was discarded. Resin was resuspended using 500 µL of HPLC-grade distilled water, incubated for five minutes, and centrifuged to discard flow-through for a total of three washes. Following washes, 40 µg of sdAb diluted in 250 µL of 100 mM MOPS buffer or the MOPS buffer alone was added to respective columns and incubated overnight. Columns were centrifuged and unbound sdAbs were collected and stored at -80°C before Western blot analysis to determine sdAb binding efficiency to the resin (see *Western Blot*). Binding of the sdAbs to the resin was also confirmed by dot blot (see *Dot Blot*). Columns were washed using 500 µL of 100 mM MOPS buffer, incubated for five minutes, and centrifuged to discard flow-through for a total of three washes. Each column then received 200 µL of ProteOn 1 M Ethanolamine hydrochloride pH 8.0 (BioRad, Cat. # 1762450) and was incubated for 4 hours to quench any remaining active site on the resin. Columns were centrifuged to discard flow-through and washed with 500 µL of 100 mM MOPS buffer 3 × 5 minutes. Each column then received 500 µg of CaSki or C33A protein lysate in a final adjusted volume of 200 µL lysis buffer and was incubated overnight. Columns were centrifuged and flow-through was collected. Columns were washed 3 × 5 minutes with 500 µL of lysis buffer without inhibitors. Proteins were then eluted using 30 µL of fresh 1X SDS loading dye with DTT and were incubated on a Belly Dancer for 30 minutes at room temperature. Columns were centrifuged for 1 minute at 1000 x g at 4°C to collect the eluted proteins. The samples were heated to 95°C for 10 minutes to denature the protein and stored at -80°C before Western Blot analysis.

### Immunofluorescence in intact cells

Cervical cancer-derived cell lines CaSki, SiHa and C33A were cultured to ∼ 70% confluency, trypsinized, and washed twice with PBS by centrifugation. Cell spots (20 000 cells resuspended in 40 µL of PBS) were prepared on Fisherbrand™ Superfrost™ Plus Microscope Slides (Fisher Scientific, Cat. # 12-550-15) using passive gravitation for 10 min. The remaining fluid was soaked up with paper towels and the spots were dried completely for 15-30 min. Hydrophobic barriers were drawn around dried cell zones using a DAKO pen (DAKO, Cat. # S2002). Cells were fixed in 4% paraformaldehyde (Fisher Scientific, Cat. # O4042500) in PBS for 10 minutes at room temperature and washed 3 × 5 minutes with PBS. Cells were permeabilized with 0.1% Triton X-100 (Cedarlane, Cat. # T8655) in PBS for 10 minutes at room temperature and washed 3 × 5 minutes with PBS. Two different methods were used for the primary antibody incubation. In the first one, sdAb at a concentration of 0.04 µg/µL in 50 µL of DAKO antibody diluent (DAKO, Cat. # M3629) was applied to cell spots and incubated overnight at 4°C as previously described (Jackson et al. 2013). Staining controls received only antibody diluent. The following day, the primary antibody was removed. The cells were immediately rinsed with chilled PBS and then washed 3 × 5 minutes with chilled PBS. A goat anti-alpaca IgG, sdAb secondary antibody with Alexa Fluor® 594 (Jackson ImmunoResearch, Cat. # 128-585-230) diluted at 1:400 in DAKO antibody diluent was then applied and incubated for 1 hour at room temperature in the dark. Samples were rinsed and washed as above. Following a final rinse in dH_2_O, 10 µL of Vectashield with DAPI (Cedarlane, Cat. # H-1000) were added to each field on the slide before adding coverslips (Fisher Scientific, Cat. # 12-544-E). For the second method, cells were blocked with 1% BSA-PBS for half an hour at room temperature and subsequent incubation of antibodies diluted in 1% BSA-PBS were performed using the same concentration as above. Fluorescent microscopy at 400X magnification was performed using a Zeiss Axiovert 200 Microscope (Carl Zeiss Canada Ltd.) equipped with an LD A-Plan 40x/0.50 Ph2 objective (Carl Zeiss Canada Ltd.) and a CCD camera with 12-bit capability (Q Imaging). Using their respective microscope filters, 10 pairs of images (1 DAPI and 1 sdAb) were taken at each spot in a pattern to cover as many areas as possible. For DAPI and sdAb images, exposures of 20 milliseconds and 1 second were used, respectively. The gain was set to 60.14% and the offset to 41.00% for all images. The images were then prepared using ImageJ (https://imagej.nih.gov/ij/): DAPI (blue) and sdAb/E6 fluorescence (red) were overlaid using the function “Color-Merge Channels”. The overlayed images were analyzed using CellProfiler (version 4.2.1) using the following pipeline: “ColorToGray”, “IdentifyPrimaryObjects (Otsu, diameter 50-120 pixel units, two-class thresholding method)”, “MeasureObjectIntensity”, “DisplayDataOnImage” and “ExportToSpreadsheet”. The DAPI image was used the identify and select the nuclei and the intensities of red fluorescence in the selected regions were measured using the function “MeasureObjectIntensity”. For each condition, 10 random fields were analyzed.

## Results and Discussion

### Review of studies developing antibodies against HPV16 E6

The development of antibodies against the E6 protein has been a long-lasting objective starting in the 1990s. During this time, many antibodies aimed at abolishing E6’s functions have been raised against the HPV16 E6 protein almost entirely for therapeutic purposes. Recently, anti-E6 antibodies have been reviewed focusing on their formats, therapeutic proprieties, and delivery methods (Donà et al. 2021). Although some antibodies have shown therapeutic potential, the search for more efficient molecules with higher therapeutic capacity is still ongoing. Here, we review in detail how these Abs have been generated and selected (Table S1) to identify potential improvements in anti-E6 Ab development. Abs are also powerful molecular biology tools used in research for various applications including Western blot, co-immunoprecipitation (Co-IP), and immunofluorescence (IF). Surprisingly, very few of the anti-E6 antibodies generated have been evaluated for their capacity to detect E6 isolated from HPV16-positive cells in Western blot or Co-IP, and, until recently, none were investigated for their E6 immunostaining capacity using fluorescence detection in natural HPV16-positive cells (Table S2).

Over the years, many conventional antibodies against the E6 protein have been reported in the literature. Such antibodies have been successfully used for targeting extracellular antigens, however their utilization for targeting intracellular proteins remains a challenge for researchers worldwide (Trenevska et al. 2017). In addition to the challenges associated with intracellular antibody delivery, their size (∼ 150 kDa) implies that they cannot enter the nucleus by passive diffusion whereas smaller molecules such as scFvs (∼ 25 kDa) may freely enter the nucleus (Wang et al. 2007). Nevertheless, several mAbs specific for the N-terminal (1F1, 6F4, 4C6), zinc-binding domain (1F5, 3B8, 3F8), or unknown epitope (F127-6G6) of the HPV16 E6 protein have been isolated from immunized BALB/c mice (Giovane et al. 1999, Masson et al. 2003, Lagrange et al. 2005, and discussed in Togtema et al. 2019). Two approaches have been used to deliver such mAbs to HPV16-positive cells, namely cationic lipid-based transfection (HiPerFect, Qiagen) and sonoporation. Using the former, 6F4, 4C6, and 3F8 were delivered into CaSki and SiHa cells. Only 4C6 was able to restore p53, albeit without apoptosis. However, HPV16-positive cell proliferation was reduced by 50% compared to cells transfected with HiPerFect alone (Courtête et al. 2007). Our lab also used HiPerFect to evaluate the therapeutic potential of 4C6 and F127-6G6 in CaSki and SiHa cells, however only F127-6G6 showed significant increases in the percentage of p53-positive cells compared to all controls. Additionally, the p53 intensity per cell was also significantly increased under these conditions, but no cytopathic effect of apoptosis was noted. For successful delivery of conventional antibodies using sonoporation, microbubbles are incubated with the mAb of interest, added to target cells, and exposed to high intensity focused ultrasound (HIFU). This causes the microbubbles to explode, creating pores in the surrounding cells, allowing the internalization of the mAbs (Togtema et al. 2012). Sonoporation delivery of F127-6G6 to CaSki and SiHa cells led to a significant increase in the percentage of p53 positive CaSki but not SiHa cells compared to sonoporation alone. Nevertheless, p53 intensity between stained cells was not significantly increased and only a non-significant proportion of apoptotic cells was observed (Togtema et al. 2012). Taken together, these data indicate that mAbs targeting the N- or C-terminal region of E6 may reduce cell proliferation, increase p53 levels, but do not induce apoptosis.

Other studies have investigated the use of smaller molecules - namely single-chain variable fragment antibodies (scFvs) derived from mAbs (Giovane et al. 1999, Sibler et al. 2003, Lagrange et al. 2007). ScFvs are composed of the variable domains of conventional antibodies’ heavy and light chains, with the two variable regions held together via a peptide linker of variable length. Typically, such molecules are expressed intracellularly via cDNA cell transfer. (Sarker et al. 2019, Kang and Seong 2020). Several scFvs have been developed using the variable domains of previously described HPV16 E6 monoclonal antibodies (scFv1F1, scFv6F4) and through mixing the heavy and light chain variable domains of such monoclonal antibodies (6F4 and 1F1 to scFv1F4 and scFv6F1) (Giovane et al. 1999, Sibler et al. 2003, Lagrange et al. 2007). To increase the solubility of the scFv1F4, random mutations were inserted to generate a solubility-enhanced antibody called 1F4-P41L (Sibler et al. 2003). Low-level apoptosis was observed in CaSki and SiHa cells transduced with scFv1F4 and 1F4-P41L (Lagrange et al. 2007). Surprisingly, apoptosis induction did not correlate with an increase in p53 levels, and the authors suggested the presence of potential p53-independent apoptotic mechanisms (Lagrange et al. 2007). Nevertheless, 1F4-P41L aggregation was still abundant (only 10% soluble), leading to toxic effects in the HPV16 E6 positive or negative cell lines tested (Lagrange et al. 2007). Such scFvs shortcomings were also observed by another group (Kang and Seong 2020), suggesting that the reducing environment of the cytoplasm could interfere with the formation of disulphide bonds linking the complementarity-determining regions (CDRs), therefore, leading to the formation of non-functional, aggregated intrabodies.

To overcome aggregation, scFvs can also be screened against the antigen of interest via phage libraries based on B cells of immunized animals, naïve non-immunized animals, or synthetic, randomly mutated CDRs (Ahmad et al. 2012). After producing and screening their synthetic scFv library with recombinant proteins (MBP- and GST-E6), Griffin et al. 2006 identified GTE6-1, showing binding to the N-terminal zinc-binding domain of E6. To quantify its therapeutic effect, SiHa cells were transfected with GFP-tagged GTE6-1, and probed for p53, keratin 18 cleavage by caspase 3 (indicating an early sign of apoptosis), and pyknotic nuclei (an indicator of late apoptosis). At 48 hours post-transfection, the percentage of GFP-expressing cells positive for p53, keratin 18 cleavage, or showing pyknotic nuclei were 60%, 80%, and 60% respectively (Griffin et al. 2006). Another scFv was developed by intracellular capture technology using a naïve library of intracellular antibodies allowing the expression of the scFv and antigen in yeast (Amici et al. 2016), facilitating the selection of more stable scFvs that can better withstand the lack of disulphide bond formation when expressed in the cytoplasm of targeted cells. Screening the library using E6 genes led to the identification of an scFv named I7, which was further modified using a nuclear localization signal (I7nuc) to target nuclear E6 more efficiently. When tested in SiHa, C33A and 293T cells retrovirally transfected with plasmids for E6 and I7nuc expression, there was a roughly 3-fold increase in p53 levels after 24 hours compared to cells transduced with the empty retroviral vector (Amici et al. 2016). It was also observed that I7nuc-transduced SiHa cells had a 19% reduction in proliferation at 24 hours, and a 50% reduction at 96 hours post-transduction, while no reduction in proliferation was observed in transduced C33A cells. Analysis of SiHa cells expressing I7nuc using AnnexinV-FITC/PI staining showed evidence of limited late apoptosis and abundant necrosis in treated cells compared to cells transfected with the mock plasmid. Based on PARP or DNA fragmentation readouts, no significant increase in early apoptosis was detected (Amici et al. 2016). Much research is still needed for scFvs’ clinical transition, thus leading us to investigate the potential of *Camelidae*-derived single-domain antibodies (sdAbs) (Hamers-Casterman et al. 1993).

*Camelidae* naturally express heavy chain only antibodies lacking the light chain and the heavy chain constant (CH1) domain of conventional Abs. The variable domain of such antibodies can be expressed as sdAbs retaining the full antigen-binding capacity of the parent molecules. The sdAbs (15 kDa) are smaller and more soluble than scFvs, and the disulphide bridges linking the scFvs CDRs are less commonly found in sdAbs, facilitating their soluble expression in bacteria or mammalian cells (Harmsen et al. 2000, Soetens et al. 2020). Additionally, they can bind concave epitopes allowing accessibility to antigens not targetable by other mAbs or scFvs (Harmsen et al. 2000, Soetens et al. 2020). Our lab recently generated 27 sdAbs by immunizing two llamas with different recombinant E6 proteins (Togtema et al. 2019). The first llama was immunized with the His6-GenScript E6 (His6-E6) protein, whose E6 corresponds to the cervical carcinoma-derived HPV16-positive cell line CaSki with the amino acid substitutions R10G and L83V compared to the reference sequence (GenBank Accession #: K02718.1) (Meissner 1999, Pattillo et al. 1977, Zehbe et al. 2009). The second llama was immunized with two recombinant E6 proteins, His6MBP-4C/4S E6 and His6MBP-F47R 4C/4S E6 (see *Recombinant Protein section* for details).

The selection of E6 binders was performed by two rounds of subtractive panning: depleting MBP binders using MBP alone and selecting for the His6MBP-4C/4S E6 and His6MBP-F47R 4C/4S E6 binders. All selected sdAbs have different CDR3 regions suggesting that they may bind to different E6 epitopes. Soluble sdAbs were expressed with a His6 tag and a hemagglutinin (HA) tag, however for 7 sdAbs, not enough soluble sdAbs were obtained and these molecules were not further or fully investigated (Togtema et al. 2019). We performed several experiments using recombinant MBP, His6MBP-4C/4S E6, and His6MBP-F47R 4C/4S E6 to characterize the sdAbs. Western blot under native PAGE conditions indicated that the 2A17 clone was the strongest His6MBP-4C/4S E6 and His6MBP-F47R 4C/4S E6 binder, which was confirmed using surface plasmon resonance which demonstrated an average binding affinity of approximately 50 nM. Using dot blots, we determined that 19 sdAbs preferentially bound to His6MBP-4C/4S E6 and His6MBP-F47R 4C/4S E6 while the sdAb C26 bound to both the MBP and the recombinant E6 proteins, suggesting that C26 was an MBP binder. In the current study, our molecules have been further analyzed, using untagged recombinant E6 and endogenous E6 proteins, leading us to uncover some diverging results, as described below.

Another group has developed the anti-E6 sdAb Nb9 (Zhang et al. (2021). In that study, a Bactrian camel was immunized using a small ubiquitin-like modifier (SUMO)-tagged E6 protein. The selection of E6 binders was performed using a bacterially expressed His-tagged E6 protein, leading to the identification of the Nb9 sdAb. In contrast to the sdAbs obtained in our previous study, Nb9 binds the denatured recombinant E6 protein in Western blot, indicating that the sdAb recognizes a linear epitope. The results were further confirmed in Western blot using CaSki and SiHa with C33A as a negative control. Immunolocalization of E6 using Nb9 in CaSki and SiHa cells detected E6 mainly in the nucleus, with minor cytoplasmic localization, with no signal detected in C33A. However, after intracellular expression of Nb9 in CaSki and SiHa cells, E6 and Nb9 were mostly found in the cytoplasm of the cells. The authors suggest that Nb9 was blocking E6’s re-entry into the nucleus (Zhang et al. 2021). Intracellular expression of Nb9 also led to increased p53 and p21 levels, as observed by Western blot, although no quantification was performed. Finally, CaSki and SiHa cells transfected with Nb9 for intracellular expression had reduced proliferation, while AnnexinV-PE/7-AAD staining of the cells showed increased apoptosis rates compared to un-transfected control cell lines (Zhang et al. 2021). Results obtained with Nb9 are encouraging due to its limited side effects and mechanism specificity compared to scFvs, as it appears to induce apoptosis through p53 restoration.

### Selecting anti-E6 sdAb candidates: divergent results to our initial study

#### SdAb clones bind differentially to a tag-free, recombinant E6 protein in combination with an alternative detection reagent

In our initial sdAb study (Togtema et al. 2019), molecules were labeled “A” or “2A” if originating from the llama immunized with His6-GenScript E6 and selected during the first (A) or second (2A) round of subtractive panning respectively or were labeled “C” if originating from the llama immunized with His6MBP-4C/4S E6 and His6MBP-F47R 4C/4S E6 and selected during the first round of subtractive panning (Togtema et al. 2019). Of the 20 earlier-tested sdAbs, only 18 were re-evaluated here because no more soluble material was available for clones A09 and A27. Among the 18 candidates, 14 resulted from immunization with His6-GenScript E6 (A05, A34, A37, A45, A46, A47, 2A03, 2A04, 2A10, 2A12, 2A15, 2A17, 2A51, and 2A78) and 4 from immunization with His6MBP-4C/4S E6 and His6MBP-F47R 4C/4S E6 (C11, C26, C36, and C38) (Togtema et al. 2019). In our previous study, clone 2A17 was found to be the most promising E6 binder to both recombinant E6 proteins (Togtema et al. 2019).

In this follow-up study, we tested the binding of 2A17 to native E6 in CaSki cells using co-IP under different conditions, but 2A17 failed to capture E6. We therefore re-evaluated the sdAbs in another round of dot blot experiments where we membrane-spotted the 3 previously used recombinant proteins (MBP, His6MBP-4C/4S E6, and His6MBP-F47R 4C/4S E6) (Togtema et al. 2019), as well as a commercially available tag-free, recombinant E6 protein with 6 cysteine to serine substitutions (E6 6C/6S). Additionally, we attempted detection of the sdAbs with a secondary antibody raised against their immunoglobulins rather than their HA tag.

Under the new conditions, 2A17’s binding to His6MBP-4C/4S E6 and His6MBP-F47R 4C/4S E6 was still strong, but it bound with much less strength to the new, tag-free E6 6C/6S and MBP (Figure 1). In addition, 2A17’s detection of E6 6C/6S and MBP was visible only at a higher exposure (data not shown). Although 2A17 originates from the llama immunized with His6-GenScript E6 without any MBP tags, the antibody selection was performed using ELISA and recombinant MBP-tagged E6 proteins (Togtema et al. 2019). The results from our current investigation indicated that 2A17’s epitope is likely composed of a fragment of both MBP and E6, such that the sdAb fails to interact with each protein independently. Some sdAbs seemed to bind similarly well to His6MBP-4C/4S E6, His6MBP-F47R 4C/4S E6, and E6 6C/6S, while not binding to MBP alone: A05, A37, A45, A46, A47, C38, 2A03, 2A04, and 2A78 (Figure 1). While C11, 2A12, and 2A15 also bound to these 3 recombinant E6 proteins, the signal detected was extremely low and only observed at a higher exposure (20 minutes compared to 1 minute). The remaining clones showed more disparity in their binding capacity to the different E6 proteins: C36 bound strongly to both His6MBP-4C/4S E6 and His6MBP-F47R 4C/4S E6 but showed very low affinity for E6 6C/6S (Figure 1). As C36 was produced in the llama immunized with both MBP-tagged E6 proteins, it could be hypothesized that, similarly to 2A17, C36 requires MBP to stabilize its interaction with E6. The 2A10 antibody bound similarly to both E6 6C/6S and His6MBP-4C/4S E6 proteins, with preferential binding to His6MBP-F47R 4C/4S E6 (Figure 1). Although A34 and 2A51 bound His6MBP-4C/4S E6 and His6MBP-F47R 4C/4S E6 in previous dot blot experiments (Togtema et al. 2019), they did not do so with any of the proteins in the current dot blots (Figure 1). This could indicate that a higher concentration of A34 and 2A51 would be needed to detect recombinant E6. It is also possible that the secondary anti-sdAb antibody used to detect the sdAbs in the current dot blots was not efficient at detecting A34 or 2A51. Nevertheless, due to other positive E6 6C/6S binders, we did not further test either hypothesis for these clones.

**Figure 1.**
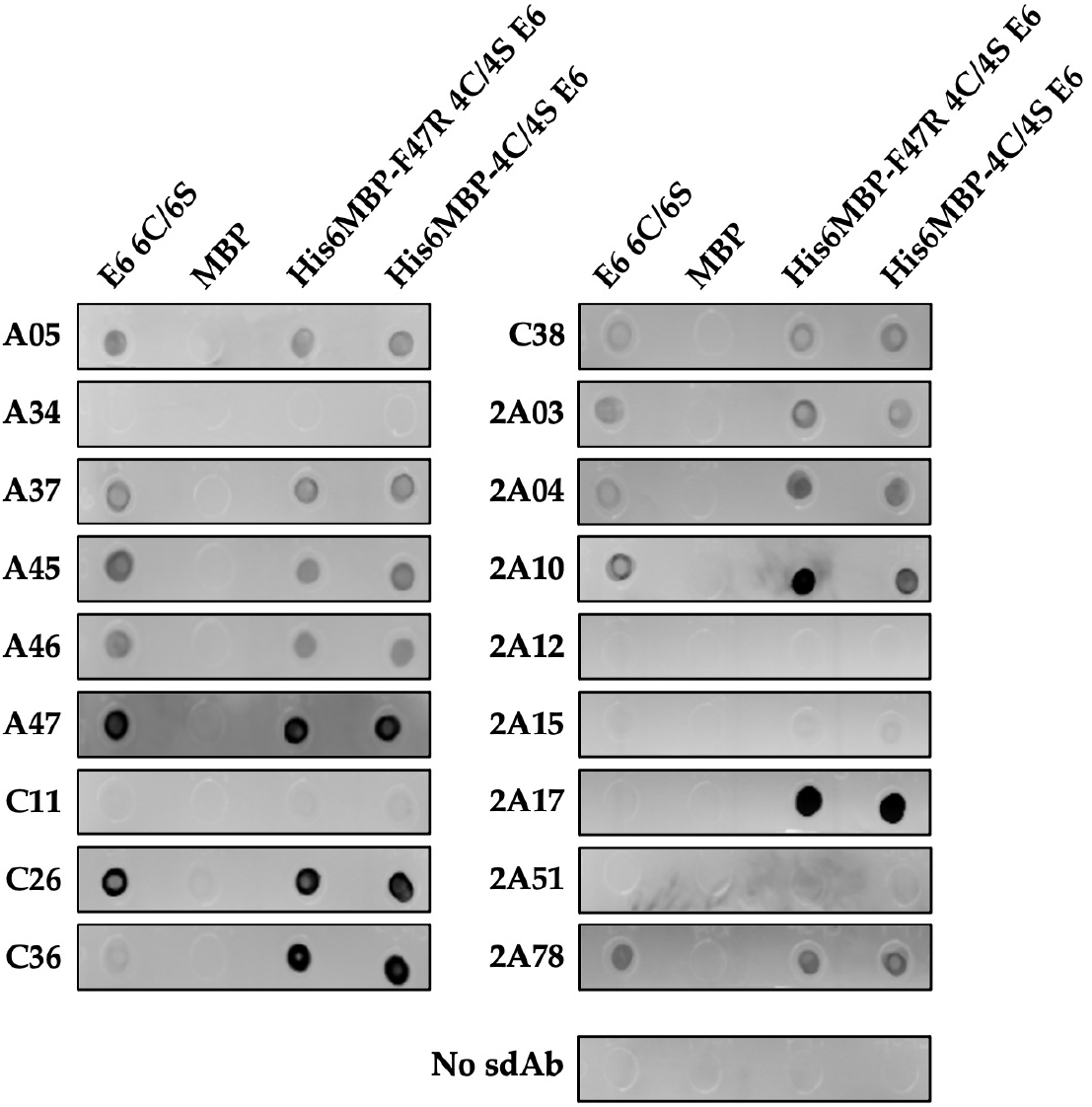
Dot blot analysis of the sdAbs’ binding capacity to different recombinant proteins. For the dot blot, 2 µg of each recombinant protein: E6 6C/6S, MBP, His6MBP-F47R 4C/4S E6, and His6MBP-4C/4S E6, was spotted on the membrane and detected using 5.4 µg/mL of sdAb. To exclude unspecific binding of the secondary anti-sdAb antibody, a dot blot was performed in the absence of sdAb [No sdAb].

In our previous publication, dot blot as well as Western blot under reducing SDS-PAGE and native PAGE conditions led to the conclusion that C26 was an MBP binder recognizing a linear epitope (Togtema et al. 2019). However, in light of our newly collected data, we could not fully exclude C26 also binding to E6. In the current dot blots with the addition of the new, tag-free E6 6C/6S, C26 seemed to preferentially bind to that recombinant E6 protein with almost no binding to MBP. Interestingly, previous ELISAs performed using His6MBP-4C/4S E6, His6MBP-F47R 4C/4S E6, and MBP alone seem to indicate an increased binding affinity to the recombinant E6 protein compared to MBP (Figure Supplemental ELISA). However, limited attention was given to these findings due to the number of MBP-E6 binders originally identified. SdAbs contain 3 CDR regions (paratopes) involved in antigen interaction. However, 30% of sdAb paratopes are composed of only one or two CDRs, while this is the case for only 10% of conventional Abs (Mitchell and Colwell 2018). Furthermore, CDR3 is the main region in sdAb affinity and specificity to their target. Since the loop containing CDR3 is also longer, CDR3 is the antibody region involved in the binding of a folded epitope (Mitchell and Colwell 2018). We hypothesized that C26 interacts with E6 and MBP proteins using different CDRs, such that C26 binds a folded epitope on E6 with higher affinity via CDR3 and a linear MBP epitope via CDR1, 2, or both. To test this hypothesis, we performed Western blot under reducing SDS-PAGE conditions to evaluate the binding of C26 to His6MBP-4C/4S E6, His6MBP-F47R E6, E6 6C/6S, and MBP (Figure 2). As anticipated, C26 reacted with these 3 initially used recombinant proteins as previously reported (Togtema et al. 2019), but not with the new E6 6C/6S supporting our hypothesis.

**Figure 2.**
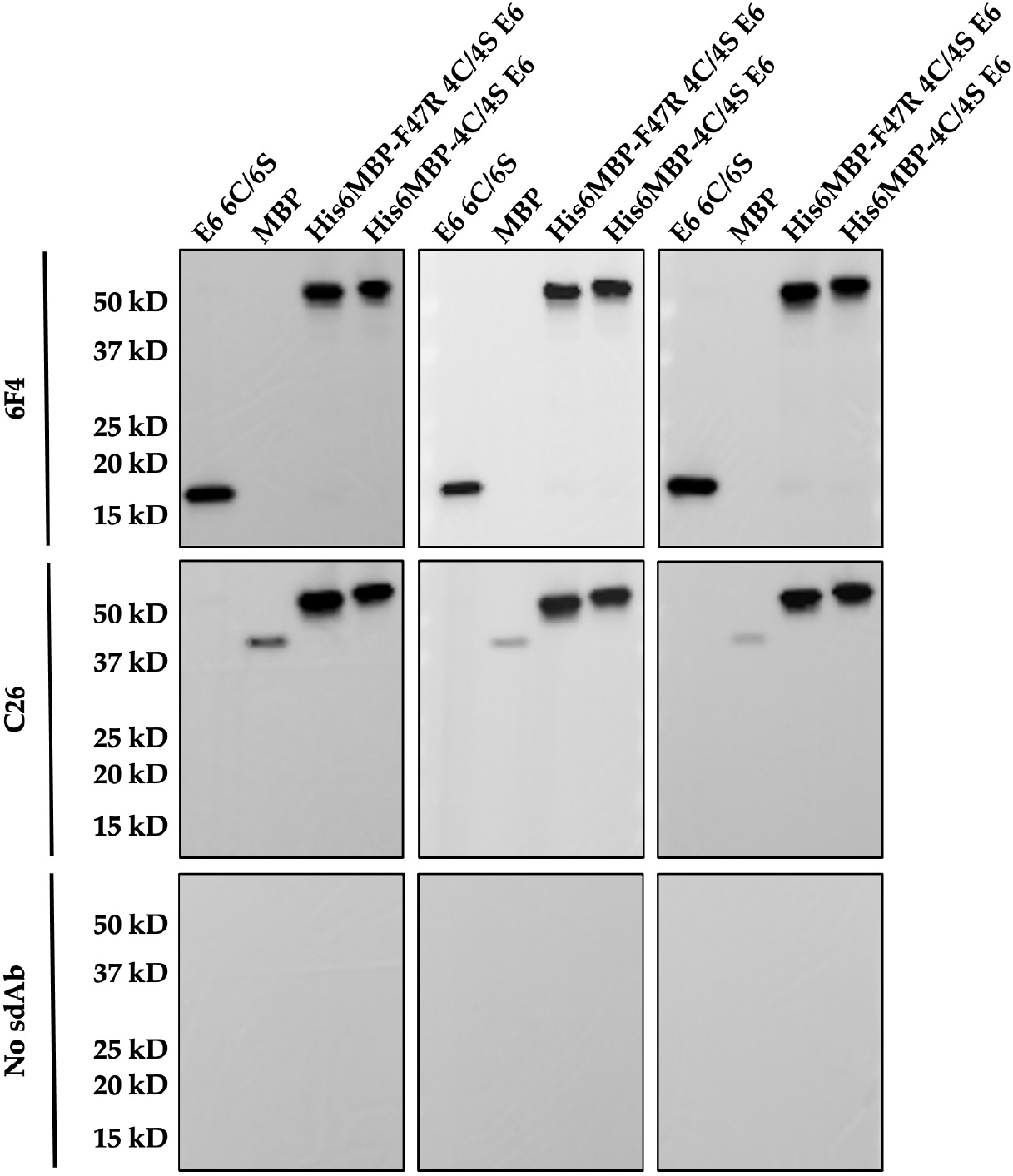
Western blots under reducing SDS-PAGE conditions. The experiments were performed in technical triplicate using 10 ng of recombinant protein: E6 6C/6S, MBP, His6MBP-F47R 4C/4S E6, and His6MBP-4C/4S E6, was prepared and denatured for each lane, respectively. Membranes were incubated with anti-E6 6F4 mAb, C26 sdAb, or secondary antibody only [No sdAb], respectively. This was followed by 1:1000 goat anti-mouse antibody conjugated to HRP for 6F4, and 1:20000 MonoRab™ Rabbit Anti-Camelid VHH Antibody conjugated to HRP for C26 and secondary antibody alone.

Following the re-evaluation of our sdAbs using the tag-free, recombinant E6 protein and the sdAb immunoglobulin-specific secondary antibody, we identified 12 sdAbs binding to the untagged E6 protein (A05, A37, A45, A46, A47, C26, C36, C38, 2A03, 2A04, 2A10, and 2A78) while the remaining antibodies (A34, C11, 2A12, 2A15, 2A17, 2A51) had either very low binding affinity to the tag-free, recombinant E6 protein or no binding at all. Finally, the dot blot results for the remaining E6 binders, A05, A37, A46, C26, 2A03 and 2A04 were confirmed in triplicate (Supplemental Dot Blot), to enable the evaluation of these sdAbs’ binding capacity to endogenous E6 - the next step in further characterization. Due to a low amount of A47 and 2A10, these sdAbs were not further evaluated in dot blot, but their binding capacity to the endogenous E6 protein was evaluated. However, for C38, A45, and 2A78, the amount was too low, and experiments were discontinued.

### Detection of sdAb binding to the endogenous E6 protein in cell lysates and intact cells

#### C26 and A37 significantly immunoprecipitate endogenous E6 in CaSki cells

E6 co-immunoprecipitation was performed using CaSki and C33A cells. Due to the low amount of endogenous E6 in SiHa cells, and the low availability of some clones, we did not attempt co-immunoprecipitating SiHa E6 using our sdAbs. Considering our sdAbs were expressed with an HA tag, we first used anti-HA magnetic beads to immobilize the antibodies, following a protocol we recently described for the immunoprecipitation of HA-E6 expressed in PHFK cells (Dayer et al. 2020). The only differences with the published interactome study methodologies were the cell type used (CaSki and C33A), the volume of magnetic beads (20 µL), and the amount of protein lysate added (1 mg). However, using this method, none of the sdAbs tested were capable of immunoprecipitating E6 (data not shown). The magnetic beads used in our experiments are coated with anti-HA mAbs which have a size of about 150 kDa. Considering this is roughly 10 times bigger than the sdAbs or E6, we speculated that the binding of the anti-HA mAb to the sdAbs likely impaired the capacity of the sdAbs to interact with E6. Instead, we used Affi-Gel resin to bind and immobilize our sdAbs. Affi-Gel contains a reactive N-hydroxy-succinimide group on a 10-atom spacer arm which covalently binds primary amines, linking the target protein to the resin. Since sdAbs contain primary amines on their amino-terminal and lysine side chain, each sdAb can be linked to the resin without the need of an anti-HA mAb. For our experiments, we used Affi-Gel 10 to immobilize basic sdAbs (pI above 6.5) and Affi-Gel 15 for acidic sdAbs (pI below 6.5), as recommended by the supplier (see *Materials and Methods* for details). Following binding of the sdAbs to the resin, we performed Western blot to confirm successful immobilization of the antibody (input compared to output of sdAb). Binding was confirmed for all sdAbs (Figure 3.A, Supplemental Co-IP 1A, 2A, 3A, 5A), but remained poor for A05 (Supplemental Co-IP 4A).

**Figure 3.**
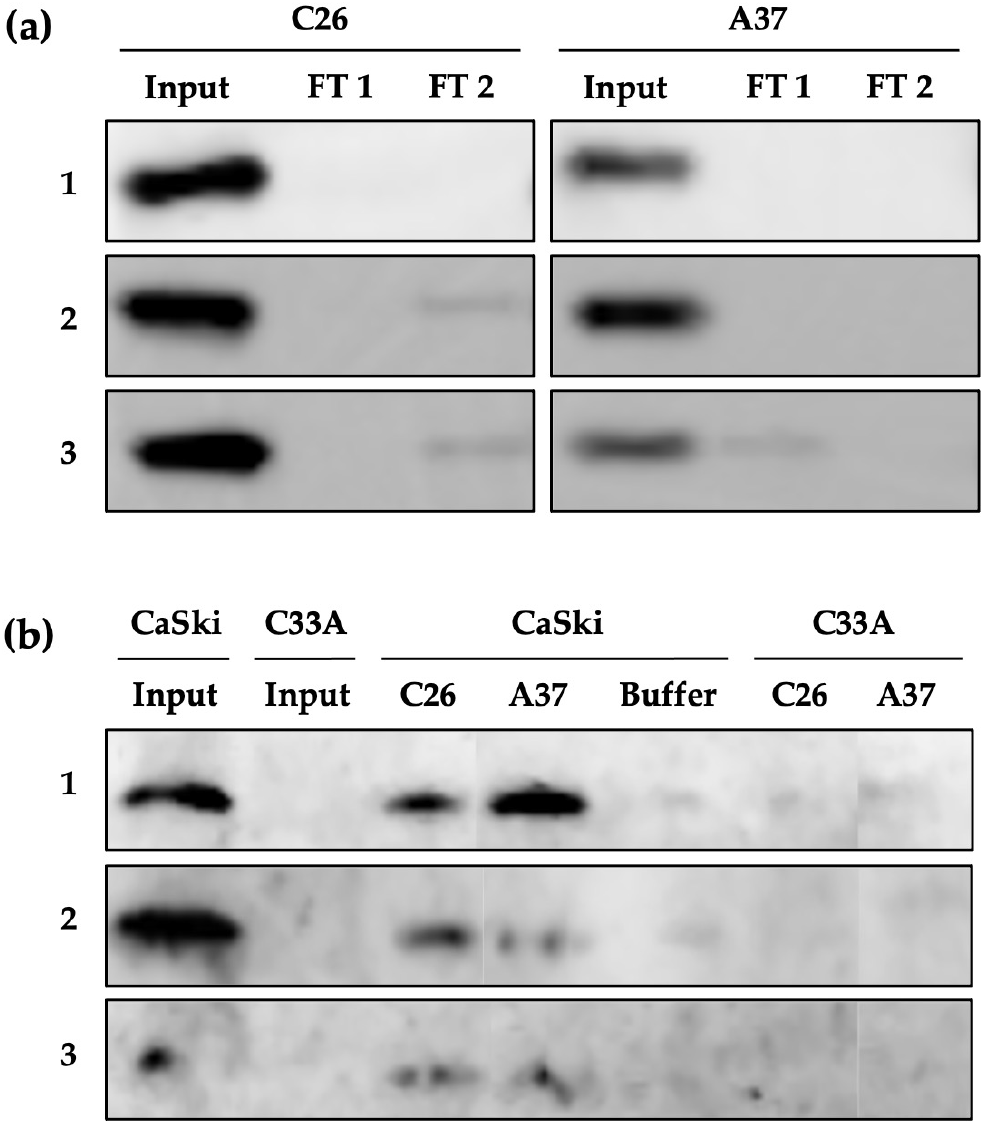
Co-IP of endogenous E6 by C26 and A37. (**a)** Confirmation of C26 and A37 binding to Affi-Gel 10 resin. Input corresponds to the 40 mg sdAb sample loaded on the resin. Flowthrough FT 1 and FT 2 correspond to the unbound antibody after incubation. The flowthrough FT1 and FT2 resin were used, respectively, with CaSki and C33A protein extract. (**b**) Co-IP results. 500 µg CaSki or C33A input samples were loaded on C26 or A37-bound resin. C26 and A37 successfully immunoprecipitated CaSki E6 in triplicate. No bands corresponding to E6’s molecular weight were observed with C26, or A37-bound resin incubated with C33A or in the resin alone control [Buffer].

E6 was immunoprecipitated with the sdAbs at varying degrees, with all trials performed in biological triplicates (3 different passages). C26 and A37 specifically immunoprecipitated CaSki E6, with no unspecific bands visible in the C33A or the CaSki + Affi-Gel 10 control samples (Figure 3). Although C26 and A37 immunoprecipitated E6 in each experiment, the efficiency of the immunoprecipitation appeared to vary. Furthermore, when comparing the CaSki input to the eluted samples (CaSki + sdAb), only a limited amount of E6 was immunoprecipitated, suggesting that further optimization was needed (Figure 3B). Candidates A05, 2A03, and 2A04 showed inconsistent results, with only a limited amount of E6 being immunoprecipitated in a single trial (Supplemental Co-IP 2, 4), suggesting that these antibodies have limited affinity to the endogenous E6 protein. A46, A47, and 2A10 did not immunoprecipitate E6 (Supplemental Co-IP 1, 3, 5), despite binding to the recombinant E6 proteins in the dot blot. This could indicate that the location of these antibody’s epitopes lies within one of the mutated regions of the recombinant E6 protein. Based on these co-IP results, C26 and A37 immunoprecipitated E6 most efficiently. Nevertheless, A05, 2A03, 2A04 were also further investigated in IF, as they did show some endogenous E6 binding capacity.

#### C26 and A37 bind endogenous E6 in intact cells, as determined by immunofluorescence, but do not raise p53 levels or induce apoptosis

Currently, there are no reliable antibodies available for the detection of E6 in intact cells using immunofluorescence (IF). Our lab previously developed a reliable method for E6 detection using the 4C6 mAb (Jackson et. al. 2013), but this method works only with enzymatic reporter molecule detection. IF allows simultaneous imaging of multiple cellular factors in the same cell and is beneficial for complementary analyses in basic and clinical HPV research. For instance, E6 localization will be essential to study its activity in intact cells and the dynamic of its interactions with host proteins. We therefore evaluated the sdAbs’ endogenous E6 binding capacity in CaSki and SiHa cells. Several steps were necessary to obtain robust results, as reported by us previously (Jackson et al. 2013): i. un-specific, nuclear, or cytoplasmic background signals were more efficiently reduced, especially in C33A cells, using a commercial antibody diluent (DAKO) than our homemade 1% BSA-PBS reagent (Supplemental IF); and ii. images were always taken with the same microscope settings and identically processed with ImageJ and CellProfiler (see *Materials and Methods* for details). A specific, nuclear fluorescence signal, albeit with differing intensity, was then observed in CaSki and SiHa cells with all sdAbs tested, with the best candidates being clones C26 and A37 (Figure 4). The average E6 signal intensity in CaSki stained with C26 was 1.09 times higher than in CaSki stained with A37, and 5.79 times higher than in SiHa stained with C26. Finally, CaSki stained with A37 were 6 times more intense than SiHa stained with A37. Together, these data confirm earlier data that E6 is mainly nuclear and that E6 expression levels are dependent on HPV copy number and are comparable with colourimetric detection for E6 immunolocalization in CaSki and SiHa cells using the 4C6 mAb, as previously reported (Jackson et al. 2013).

**Figure 4.**
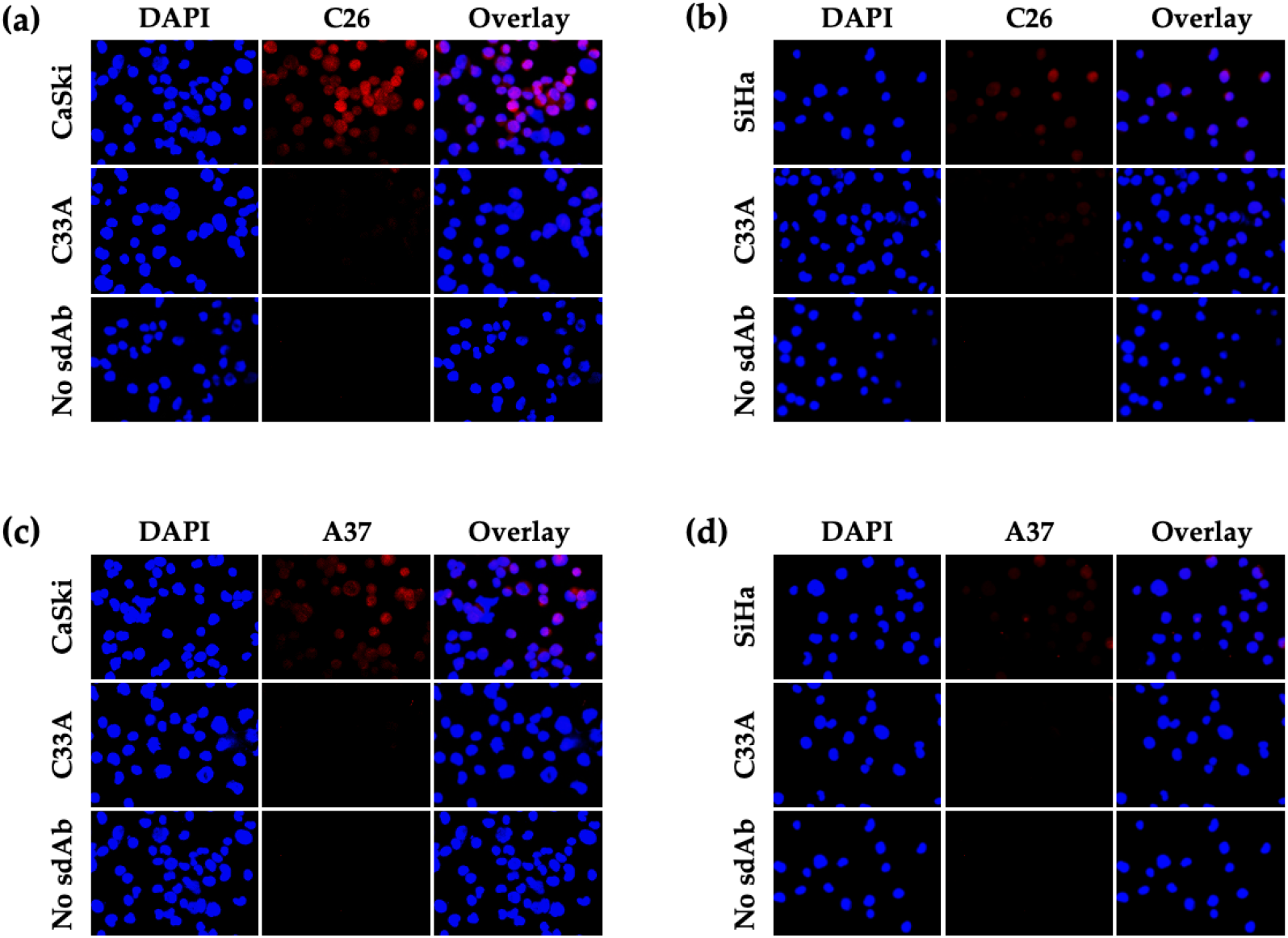
Immunodetection of E6 in CaSki and SiHa cells using C26 and A37. (**a**) and (**b**) C26 detection of E6 in CaSki and SiHa cells, respectively. (**c**) and (**d**) A37 detection of E6 in CaSki and SiHa cells, respectively. The HPV16-negative cell C33A was used as a control in each experiment. To detect unspecific binding of the secondary antibody alone, similar experiments were performed in parallel on CaSki and SiHa cells without the addition of sdAb [No sdAb].

Positive nuclear signals were also observed using 2A03 and 2A04 in CaSki and SiHa, but overall, the signal was less intense compared to C26 or A37. This is in support of 2A03 and 2A04’s lower binding affinity to the endogenous E6 protein, as observed in co-IP (Supplemental Co-IP 2A). In contrast to C26 and A37, weak background signals were consistently detected in the C33A control (Supplemental IF). A low level of background signal in HPV16-negative cell lines was also observed in previous studies with 6F4 and 4C6 (Masson et al. 2003, Jackson et al. 2013). Finally, A05 did not yield any signal in CaSki, SiHa, or C33A when using the DAKO reagents. However, A05 detected E6 in the nucleus in one experiment using 1% BSA-PBS (Supplemental IF). Because C26 and A37 showed the strongest IF signals evidencing binding to endogenous E6, we also checked if p53 levels were raised and apoptosis was induced.

Antibodies blocking the E6 protein may restore p53 and potentially induce apoptosis (Togtema et al. 2012 and references herein). To evaluate such an effect as a first proxy for the therapeutic potential of the sdAbs under study, we performed IF post C26 mRNA and A37 mRNA transfection to evaluate if p53 or PARP-1 levels differed from transfected to non-transfected cells. Treatment via C26 mRNA and A37 mRNA transfection lasted for 48 hours, however no relevant effects on p53 restoration, PARP-1 activation, or any morphological signs for apoptosis were observed in the treated cells visually by the authors, nor using CellProfiler (Supplemental IF p53 PARP). This is in partial contrast to our study using full-sized Abs against HPV16 E6, where we showed raised p53 levels in treated *vs*. untreated cells (Togtema et al. 2012).

## Conclusion

In the past two decades, various types of antibody molecules targeting the E6 protein such as conventional antibodies, scFvs, and sdAbs have been developed. However, none have achieved clinical transition. In this study, we have reviewed current literature as well as further evaluated previously isolated sdAbs against the native E6 protein. In addition, we report the first use of mRNA delivery with anti-E6 sdAbs and present a reliable methodology for this technique. Using this methodology, we achieved efficient delivery of these sdAbs into CaSki, SiHa and C33A cells, and noted their localization mainly in the nuclei of the tested target cells. We hypothesize that the sdAb candidate C26 binds a folded E6 epitope through its CDR3, and a linear MBP epitope through its CDR1 or 2 or both. This dual C26 property is most likely due to the antibody selection process using MBP-tagged E6. Our current data indicates that the presence of a tag during the immunization and/or antibody selection stage strongly interferes with the specificity of the antibodies selected. Thus, future research into the generation of anti-E6 antibodies should discontinue the use of tagged proteins, and instead prioritize the use of untagged reagents, such as the commercially available E6 6C/6S molecule.

While C26 and A37 showed a reproducible endogenous E6 binding ability in co-IP and IF experiments, the co-IP results for both sdAbs remained low. This could indicate that these candidates have limited binding affinity to the endogenous E6 protein. Based on our results with C26, such sdAbs likely bind the folded E6 protein but E6 is also known to interact with 50 or more host proteins (summarized in Dayer et al. 2020 with references therein) using residues distributed throughout the protein. This means that any molecule targeting E6 must compete with host proteins. Next steps should therefore include the characterization of paratopes, and epitopes involved in the sdAb-E6 interactions. This study presents a pipeline for evaluating anti-E6 antibody molecules for various applications in the clinic and in research. Considering the limited availability of E6 antibodies, especially for *in situ* detection, the molecules described herein have applications in both co-immunoprecipitation and immunofluorescence microscopy.

## Supporting information

Supplemental ELISA

Supplemental Dot Blot

Supplemental IF p53 and PARP

sdAb Protein Sequences

## Supplementary Materials

Table S1, Table S2, Figure Supplemental ELISA, Figure Supplemental Dot Blot, Figure Supplemental Co-IP 1-5, Figure Supplemental IF, Figure Supplemental IF p53 PARP, sdAb sequences.

## Author Contributions

Conceptualization, I.Z. and M.T.; methodology, G.D., A.F., I.Z.; software, G.D.; formal analysis, G.D., A.F., T.T., I.Z.; investigation, G.D., A.F., T.T., IZ; writing—original draft preparation, G.D., A.F., I.Z.; writing—review and editing, G.D., A.F., M.T, I.Z.; visualization, G.D., A.F., I.Z.; supervision, G.D., I.Z.; project administration, I.Z.; funding acquisition, I.Z. All authors have read and agreed to the published version of the manuscript.

## Funding

This research was funded by a Natural Sciences and Engineering Research Council of Canada (NSERC) grant to I.Z. (NSERC Discovery Grant #RGPIN-2015-03855), two Mathematics of Information Technology and Complex Systems grant to G.D. (Mitacs Elevate Postdoctoral Fellowship IT15879, Mitacs Business Strategy Internship IT28944), an Ontario Graduate Scholarship to A.F., and NSERC Graduate Student Scholarship to M.T. (PGS-D3 460717-2014).

## Acknowledgments

We are grateful to Drs. Jamshid Tanha and Greg Hussack (National Research Council, Canada) for overseeing the production of the sdAbs during MT’s five visits to their lab.

## Conflicts of Interest

The authors declare no conflict of interest.

## Notes

### Competing Interest Statement

The authors have declared no competing interest.

