## Supplementary material for "Targeting the Human Papillomavirus 16 E6 Oncoprotein with Antibodies": Table S1 and S2

**Table S1.** Summary of anti-HPV16 E6 antibody production and selection. (References to be added, numerically)

| Name | Type | Origin | Selection | Epitope 158 Amino Acid E6 |
|---|---|---|---|---|
| 6F4[1,2,3] | mAb | BALB/c mice immunized with GST-E6 | ELISA using MBP and GST-E6 | N-terminal residues 9-15 (FQDPQER) |
| 1F1[1,2] | mAb | BALB/c mice immunized with GST-E6 | ELISA using MBP and GST-E6 | N-terminal residues 9-15 (FQDPQER) |
| 4C6[2,3,4] | mAb | BALB/c mice immunized with GST-E6 | ELISA using MBP and GST-E6 | N-terminal residues 9-15 (FQDPQER) |
| 1F5[5] | mAb | BALB/c mice immunized with MBP-16E6 C-terminal ZBD | ELISA using 16E6 C-terminal ZBD | residues 127-132 (DKKQRF) |
| 3B8[5] | mAb | BALB/c mice immunized with MBP-16E6 C-terminal ZBD | ELISA using 16E6 C-terminal ZBD | residues 123-127 (QRHLD) |
| 3F8[3,5] | mAb | BALB/c mice immunized with MBP-16E6 C-terminal ZBD | ELISA using 16E6 C-terminal ZBD | residues 133-138 (HNIRGR) |
| F127-6G6[4] | mAb | BALB/c mice immunized with E6 C-terminal region | Unknown | Unknown |
| scFv1F1[1] | scFv | Heavy and light chain variable domains of 1F1 | — | Likely the same as 1F1 |
| scFv6F4[1] | scFv | Heavy and light chain variable domains of 6F4 | — | Likely the same as 6F4 |
| scFv1F4[1,6] | scFv | Heavy chain of 1F1 with light chain of 6F4 | — | Likely the same as 1F1 and 6F4 |
| scFv6F1[1] | scFv | Heavy chain of 6F4 and light chain of 1F1 | — | Likely the same as 1F1 and 6F4 |
| 1F4-P41L[1,6] | scFv | Solubility-enhanced scFv1F4 | — | Likely the same as 1F1 and 6F4 |
| GTE6-1[7] | scFv | scFv synthetic library | Library screened with Immuno tubes coated with MBP- or GST-E6 | Within residues 43-83 |
| I7/I7nuc[8] | scFv | Naïve library of intracellular antibodies | Intracellular selection using E6 genes | Unknown |
| Nb9[9] | sdAb | Bactrian camel immunized with SUMO-tagged E6 | Phage display library screened using E6 | Unknown linear epitope |
| C26 | sdAb | Llama immunized with MBP-E6 | Phage display library using MBP-E6 | Unknown folded epitope |
| A37 | sdAb | Llama immunized with CaSki-E6 | Phage display library using MBP-E6 | Unknown folded epitope |

**Table S2.** Summary of the evaluation of anti-E6 antibodies binding to the endogenous E6 protein and therapeutic potential. (References to be added, numerically)

| | | Western Blot | | Immunoprecipitation | | Immunofluorescence Microscopy | | Therapeutic Potential | | |
|---|---|---|---|---|---|---|---|---|---|---|
| **Name** | **Type** | **Cell / Control** | **Successful?** | **Cell / Control** | **Successful?** | **Cell / Control** | **Successful?** | **Cell / Control** | **Molecule Delivered** | **Results** |
| 6F4[1,2,3] | mAb | CaSki / C33A | Yes | CaSki, SiHa, E6 transfected COS-1 / HeLa, C33A, untransfected COS-1 | Yes | E6 transfected HeLa and HaCat cells / Empty vector-transfected HeLa and HaCat cells | Yes, mainly nuclear with some cytoplasmic localization | CaSki, SiHa | Protein | No effect |
| 1F1[1,2] | mAb | CaSki / C33A | Yes | CaSki / C33A | Yes | — | — | Not tested; to the best of our knowledge | | |
| 4C6[1,2,4] | mAb | — | No | CaSki, SiHa, E6 transfected COS-1 / HeLa, C33A, untransfected COS-1 | Yes | — | — | CaSki, SiHa | Protein | ↑ p53 levels |
| | | | | | | | | CaSki, SiHa | Protein | No effect |
| 1F5[5] | mAb | — | — | E6-transfected COS-1 cells, HeLa cells transfected with HPV16 E6, CaSki / Untransfected COS-1 and HeLa cells | Yes, but not for 1F5 with CaSki | COS-1 or HeLa transfected with either EGFP-E6 or EGFP-E6 C-terminal ZBD * | Yes, mainly nuclear with some cytoplasmic localization | Not evaluated due to failed RRLS inhibition of p53 degradation | | |
| 3B8[5] | mAb | — | No | E6-transfected COS-1 cell, HeLa cells transfected with HPV16 E6, CaSki / Untransfected COS-1 and HeLa cells | Yes | COS-1 or HeLa transfected with either EGFP-E6 or EGFP-E6 C-terminal ZBD * | Yes, mainly nuclear with some cytoplasmic localization | Not evaluated due to failed RRLS inhibition of p53 degradation | | |
| 3F8[3,5] | mAb | — | No | E6-transfected COS-1 cell, HeLa cells transfected with HPV16 E6, CaSki / Untransfected COS-1 and HeLa cells | Yes | COS-1 or HeLa transfected with either EGFP-E6 or EGFP-E6 C-terminal ZBD * | Yes, mainly nuclear with some cytoplasmic localization | CaSki, SiHa | Protein | No effect*** |
| F127-6G6[4] | mAb | Unknown | Unknown | Unknown | Unknown | Unknown | Unknown | CaSki, SiHa | Protein | p53 restoration with CaSki, not SiHa |
| scFv1F1[1] | scFv | — | No | — | No | — | — | — | — | — |
| scFv6F4[1] | scFv | — | No | — | No | — | — | — | — | — |

**Table 2. continued.**

| | | Western Blot | | Immunoprecipitation | | Immunofluorescence Microscopy | | Therapeutic Potential | | |
|---|---|---|---|---|---|---|---|---|---|---|
| **Name** | **Type** | **Cell / Control** | **Successful?** | **Cell / Control** | **Successful?** | **Cell / Control** | **Successful?** | **Cell/Control** | **Molecule Delivered** | **Results** |
| scFv1F4[1,6] | scFv | — | No | — | No | — | No | CaSki, SiHa / HeLa, A549 | DNA | ↓ proliferation in HPV16+ cells, no p53 restoration, low level apoptosis in CaSki and SiHa |
| scFv6F1[1] | scFv | — | No | — | No | — | No | — | — | — |
| 1F4-P41L[1,6] | scFv | — | No | — | No | — | No | CaSki, SiHa / HeLa, A549 | DNA | ↓ proliferation in HPV16+ cells, no p53 restoration, low level apoptosis in CaSki and SiHa |
| GTE6-1[7] | scFv | — | No | — | No | E6 transfected COS-7 cells ** | Yes, nuclear localization | SiHa / C33A, GFP-transduced SiHa | DNA | ↑ p53 expression, ↑ apoptosis |
| I7/I7nuc[8] | scFv | — | No | — | No | — | No | SiHa cells transfected with I7nuc or mock plasmid / C33A, HeLa | DNA | ↓ proliferation, limited apoptosis, abundant necrosis |
| Nb9[9] | sdAb | — | No | — | No | CaSki, SiHa / C33A | Mainly nuclear with some cytoplasmic signal | Transfected CaSki, SiHa / Untransfected CaSki, SiHa | DNA | ↓ proliferation, increased p53 levels, apoptosis |
| C26 | sdAb | CaSki, SiHa / C33A | Yes | CaSki / C33A | Yes | CaSki, SiHa / C33A | Mainly nuclear localization | CaSki, SiHa / C33A | RNA | Limited effects on cell proliferation |
| A37 | sdAb | — | No | CaSki / C33A | Yes | CaSki, SiHa / C33A | Mainly nuclear localization | CaSki, SiHa / C33A | RNA | No effect |

* 6F4 used as a positive control
** Results were compared to C1P5 or FLAG-tag E6
*** It failed in RRLS, however was tested anyways
Acronym definition: mAb (monoclonal antibody), sdAb (*Camelidae*-derived single-domain antibody also called VHH or nanobody (Nb)), scFv (single chain fragment variable antibody), ZBD (zinc-binding domain), MBP (maltose binding protein), GFP (green fluorescent protein).
