## Supplemental ELISA for "Targeting the Human Papillomavirus 16 E6 Oncoprotein with Antibodies"

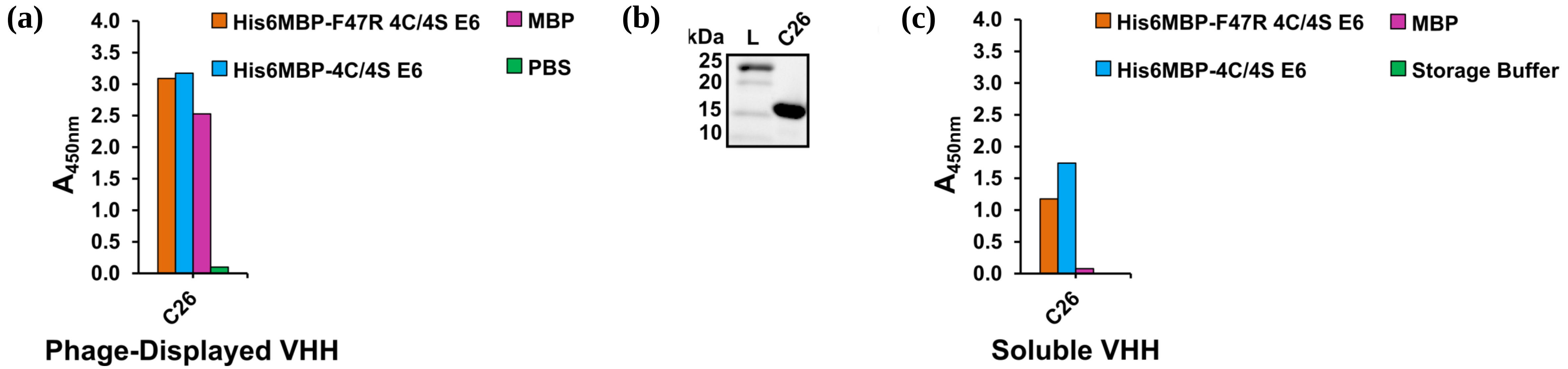


**Supplemental ELISA. (a)** Phage ELISA indicated that C26 was an MBP-binder, as phage displayed that VHH demonstrated an affinity for both His6MBP-E6 proteins as well as MBP. Clones beginning with "C" were isolated from Library #2 Round 1 eluted phage titer plates. **(b)** Reducing SDS-PAGE was used to analyze the purity of the VHH following IMAC elution. Approximately 3 µg of the sample was loaded on the gel. **(c)** An ELISA using 100 µg/mL of C26 demonstrated that this purified VHH was functional and maintained its MBP-binding characteristics.
