## Supplemental Dot Blot for "Targeting the Human Papillomavirus 16 E6 Oncoprotein with Antibodies"

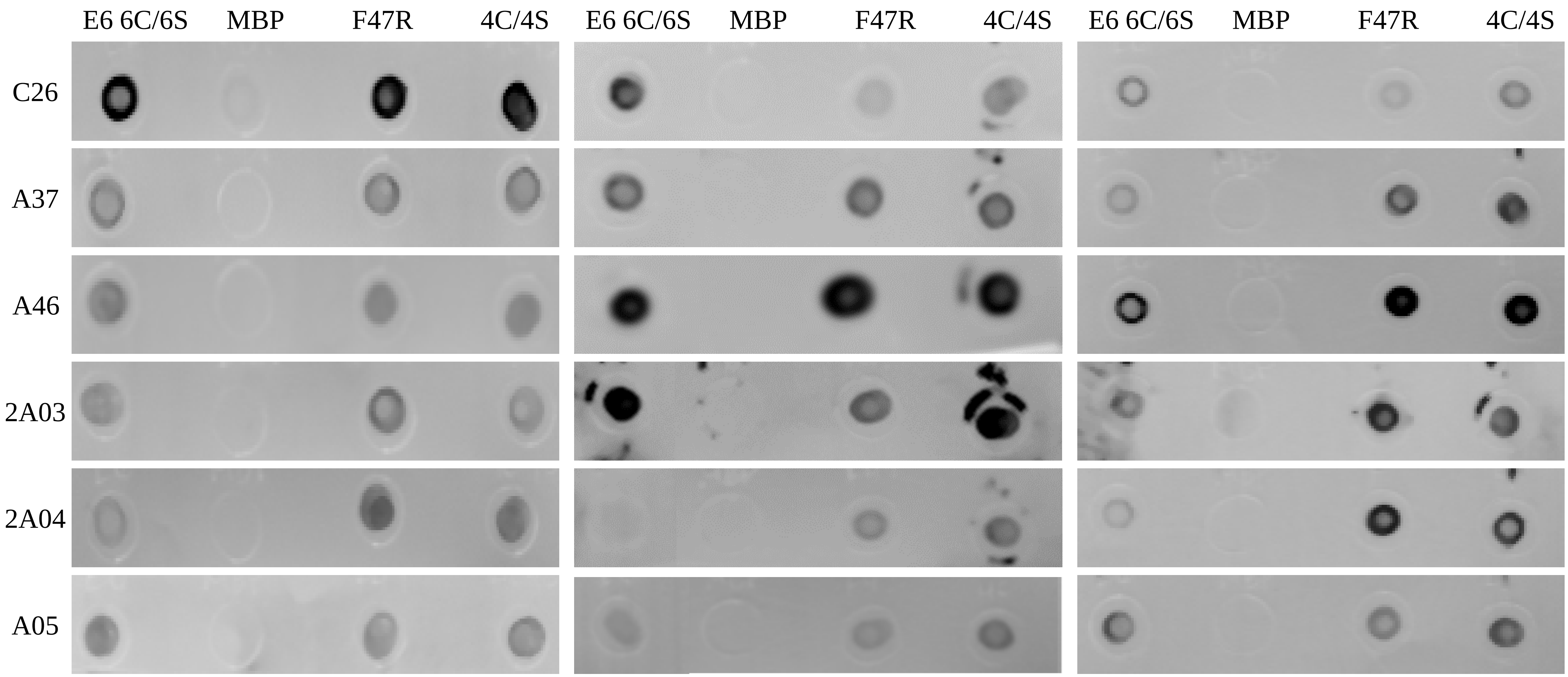


**Supplemental Dot Blot.** (Left) Dot blot results from the original dot blot experiment (Figure 1). (Middle and Right) Each spot contains 4 µL of the respective recombinant E6, E6-MBP, or MBP protein at a concentration of 0.5 µg/µl. Membranes were incubated overnight in a 5 mL blocking solution containing 5.4 µg/mL of the respective sdAb, followed by a secondary antibody incubation with MonoRab™: Rabbit Anti-Camelid VHH Antibody [HRP] in a blocking solution overnight. Membranes were visualized through chemiluminescence, UVP™ Gel Imaging System, and VisionWorks software.
