## Supplemental Co-IP for "Targeting the Human Papillomavirus 16 E6 Oncoprotein with Antibodies"

### Supplemental Co-IP: 1

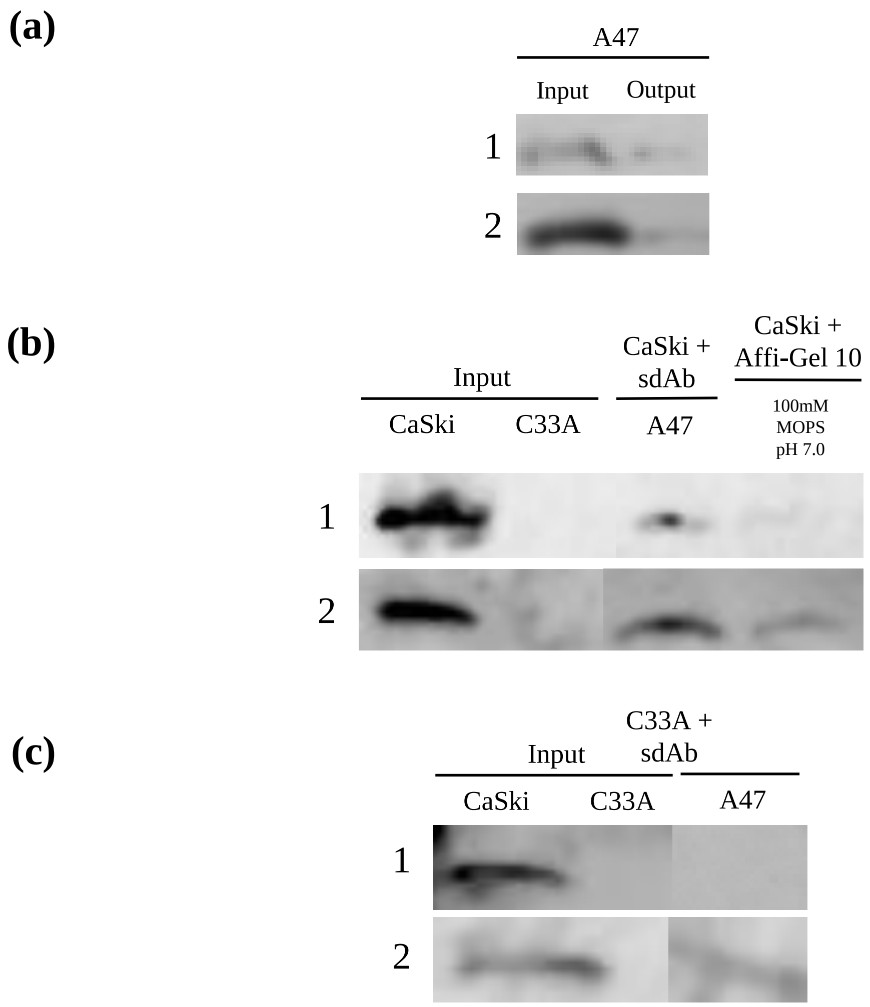


**Co-immunoprecipitation results for A47.** (a) A47 input and output for trials 1 and 2; input contains 20 μg of A47 sdAb. Membranes were visualized using HA-HRP antibody 1:1000. (b) CaSki immunoprecipitation and resin alone control results for trials 1 and 2. Membranes were visualized using 2.69 μg/μL 6F4 anti-E6 antibody. (c) C33A control immunoprecipitation results for trials 1 and 2. Membranes visualized using 2.69 μg/uL 6F4 anti-E6 antibody. 500 μg of protein lysate for both CaSki and C33A were used in both trials.

### Supplemental Co-IP: 2

(a)

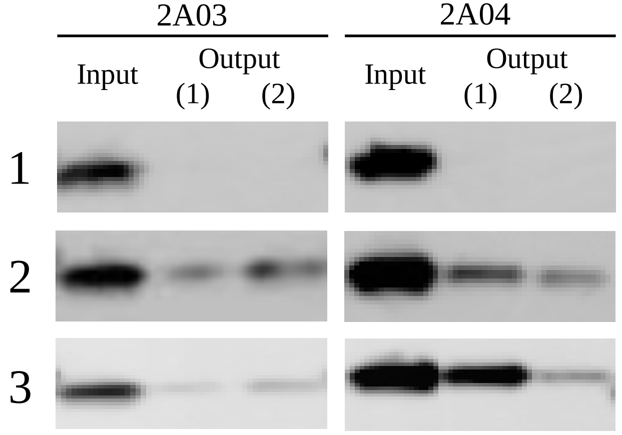


(b)

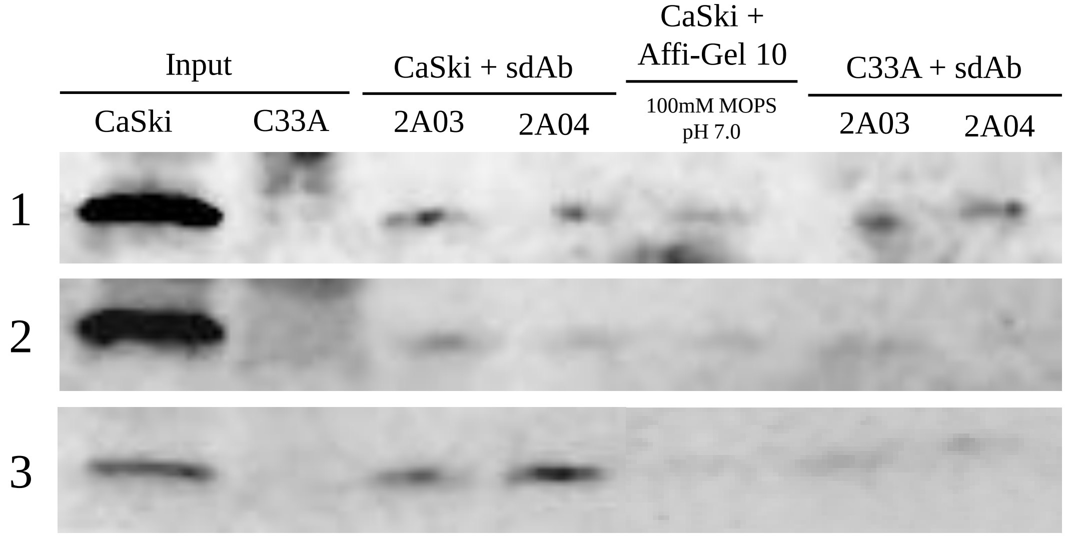


**Co-immunoprecipitation results for 2A03 and 2A04.** (a) 2A03 and 2A04 input and output for trials 1-3; input contains 40 μg of respective sdAbs. Membranes were visualized using HA-HRP antibody 1:1000. (b) CaSki and C33A immunoprecipitation and resin alone control results for trials 1-3. Membranes were visualized using 2.69 μg/uL 6F4 anti-E6 antibody. 500 μg of protein lysate for both CaSki and C33A were used in both trials.

### Supplemental Co-IP: 3

(a)

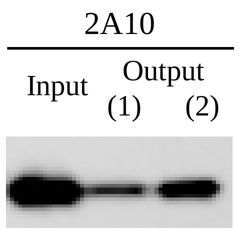


(b)

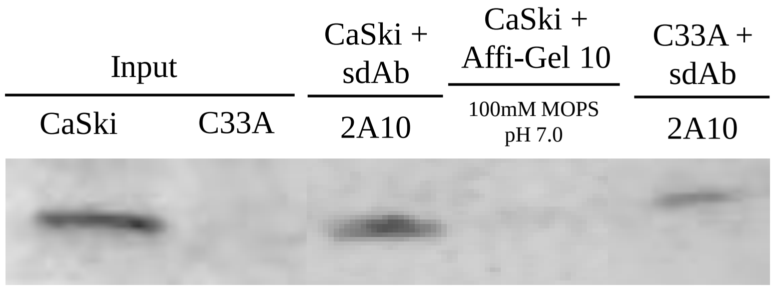


**Co-immunoprecipitation results for 2A10.** (a) 2A10 input and output; input contains 40 μg of 2A10. Membranes were visualized using HA-HRP antibody 1:1000. (b) CaSki and C33A immunoprecipitation and resin alone control results. Membranes were visualized using 2.69 μg/uL 6F4 anti-E6 antibody. 500 μg of protein lysate for both CaSki and C33A were used in both trials.

### Supplemental Co-IP: 4

**(a)**

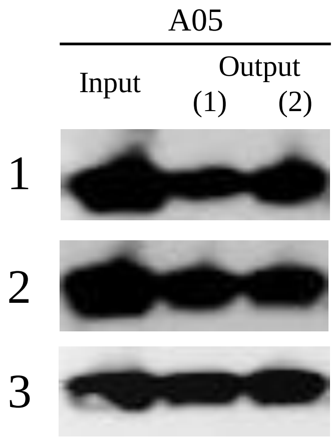


**(b)**

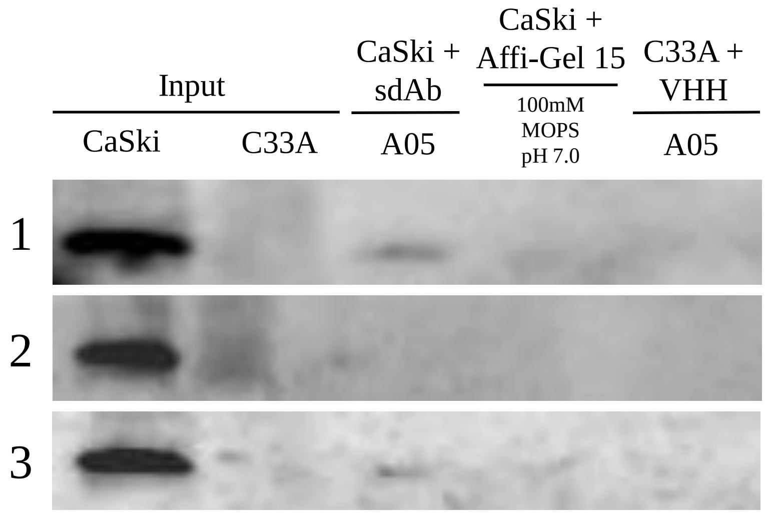


**Co-immunoprecipitation results for A05.** (a) A05 input and output for trials 1-3; input contains 40 μg of A05 sdAb. Membranes were visualized using HA-HRP antibody 1:1000. (b) CaSki and C33A immunoprecipitation and resin alone control results for trials 1-3. Membranes were visualized using 2.69 μg/uL 6F4 anti-E6 antibody. 500 μg of protein lysate for both CaSki and C33A were used in both trials.

### Supplemental Co-IP: 5

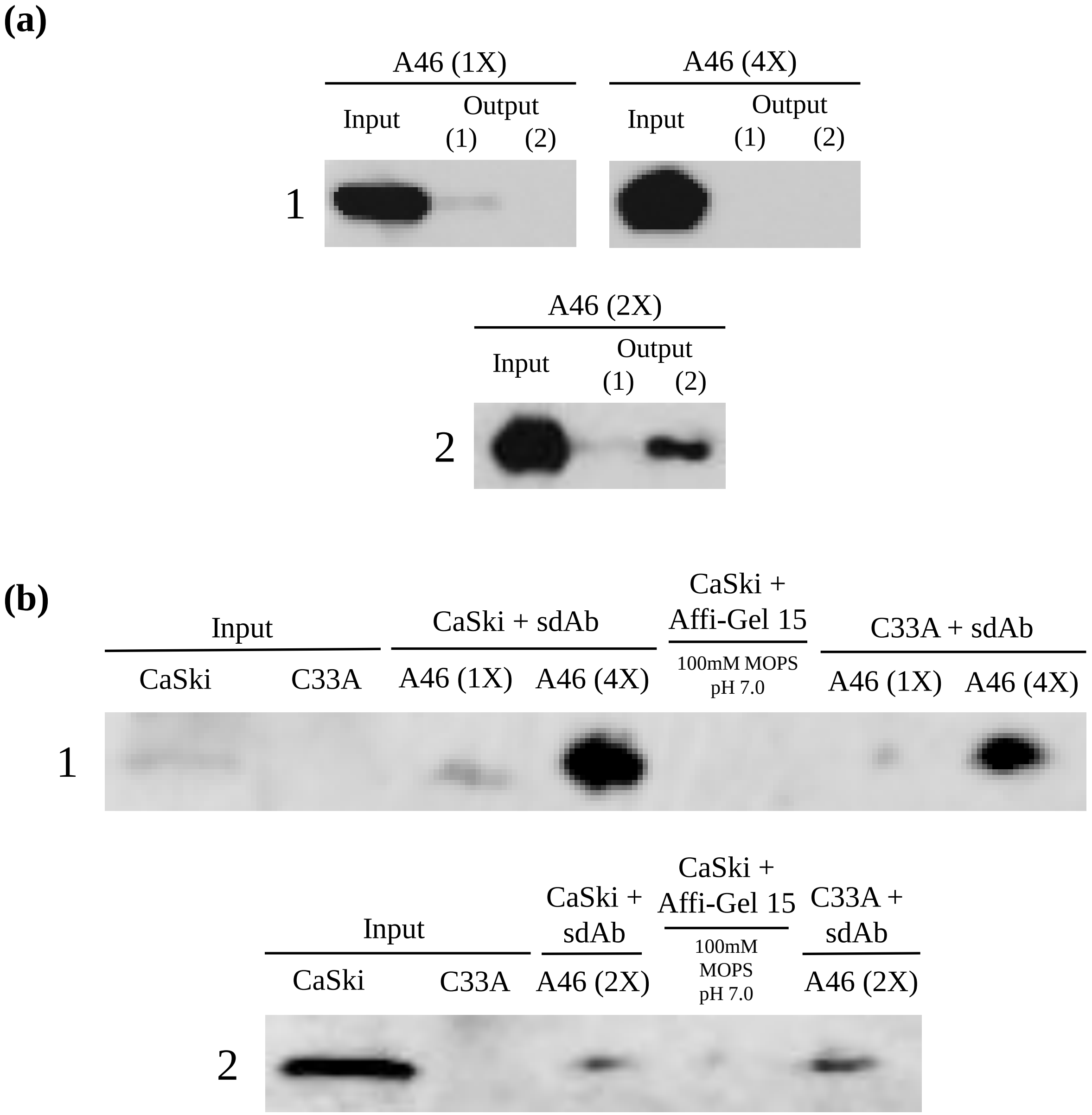


**Co-immunoprecipitation results for A46.** (a) A46 input and output for trials 1 and 2. Due to the non-specific signal observed in the control using A46, different amounts of sdAbs were added to the resin. A46 1X input contains 10 μg of sdAb, A46 2X contains 20 μg of sdAb, and A46 4X contains 40 μg of sdAb. Membranes were visualized using HA-HRP antibody 1:1000. (b) CaSki and C33A immunoprecipitation and resin alone control results for trials 1-2. Membranes were visualized using 2.69 μg/μL 6F4 anti-E6 antibody. 500 μg of protein lysate for both CaSki and C33A were used in both trials.
