## Supplementary material for "Targeting the Human Papillomavirus 16 E6 Oncoprotein with Antibodies": Supplemtal IF

### Supplemental IF: C26

**CaSki C26 (BSA-PBS)**

DAPI sdAb Overlay

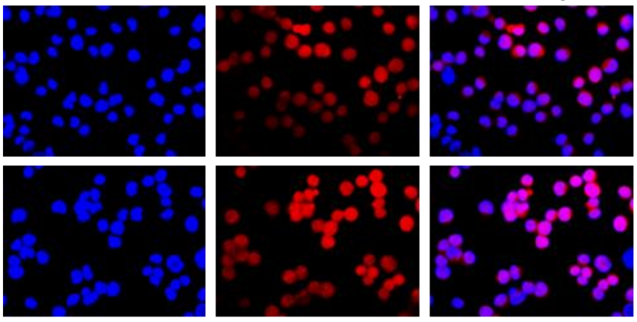

**C33A C26 (BSA-PBS)**

DAPI sdAb Overlay

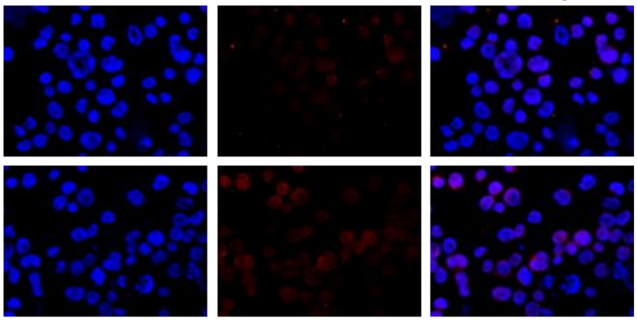

**CaSki C26 (Antibody Diluent)**

DAPI sdAb Overlay

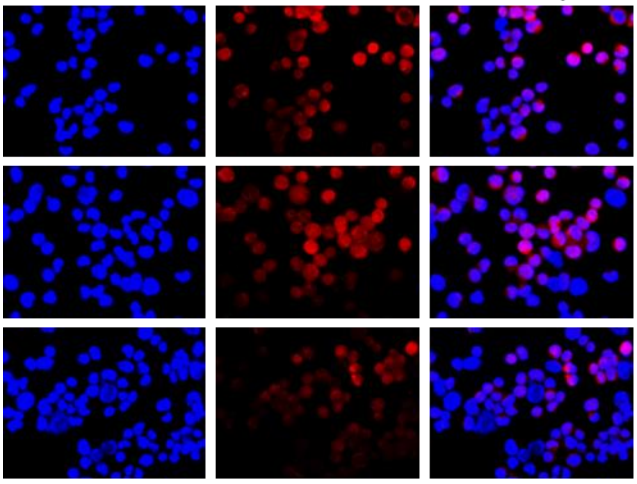

**C33A C26 (Antibody Diluent)**

DAPI sdAb Overlay

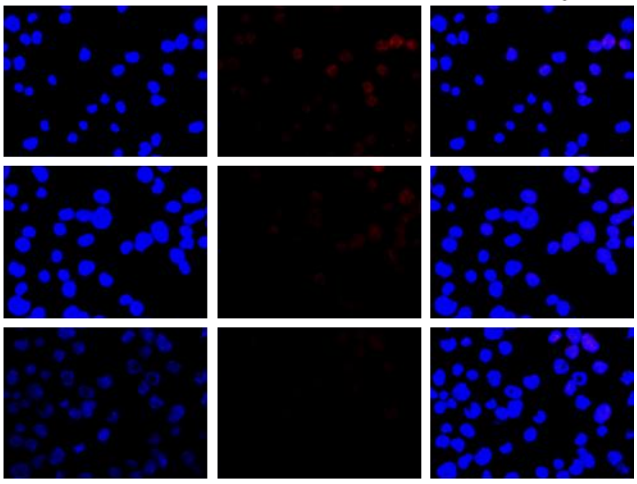

**SiHa C26 (Antibody Diluent)**

DAPI sdAb Overlay

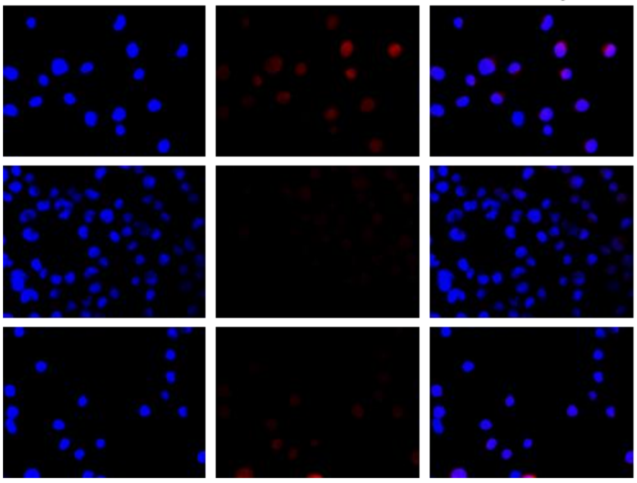

**C33A C26 (Antibody Diluent)**

DAPI sdAb Overlay

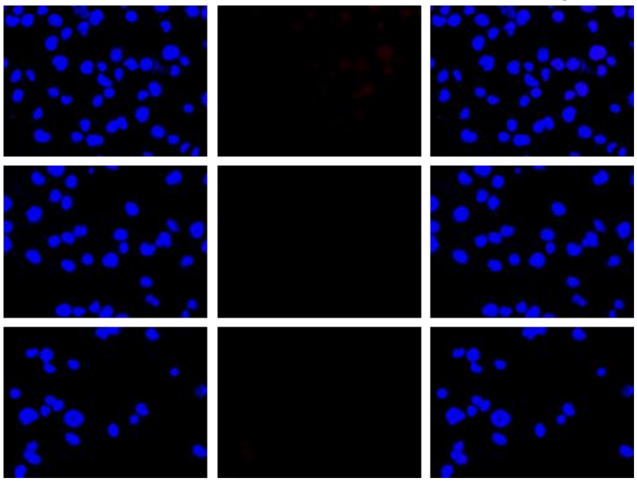

### Supplemental IF: A37

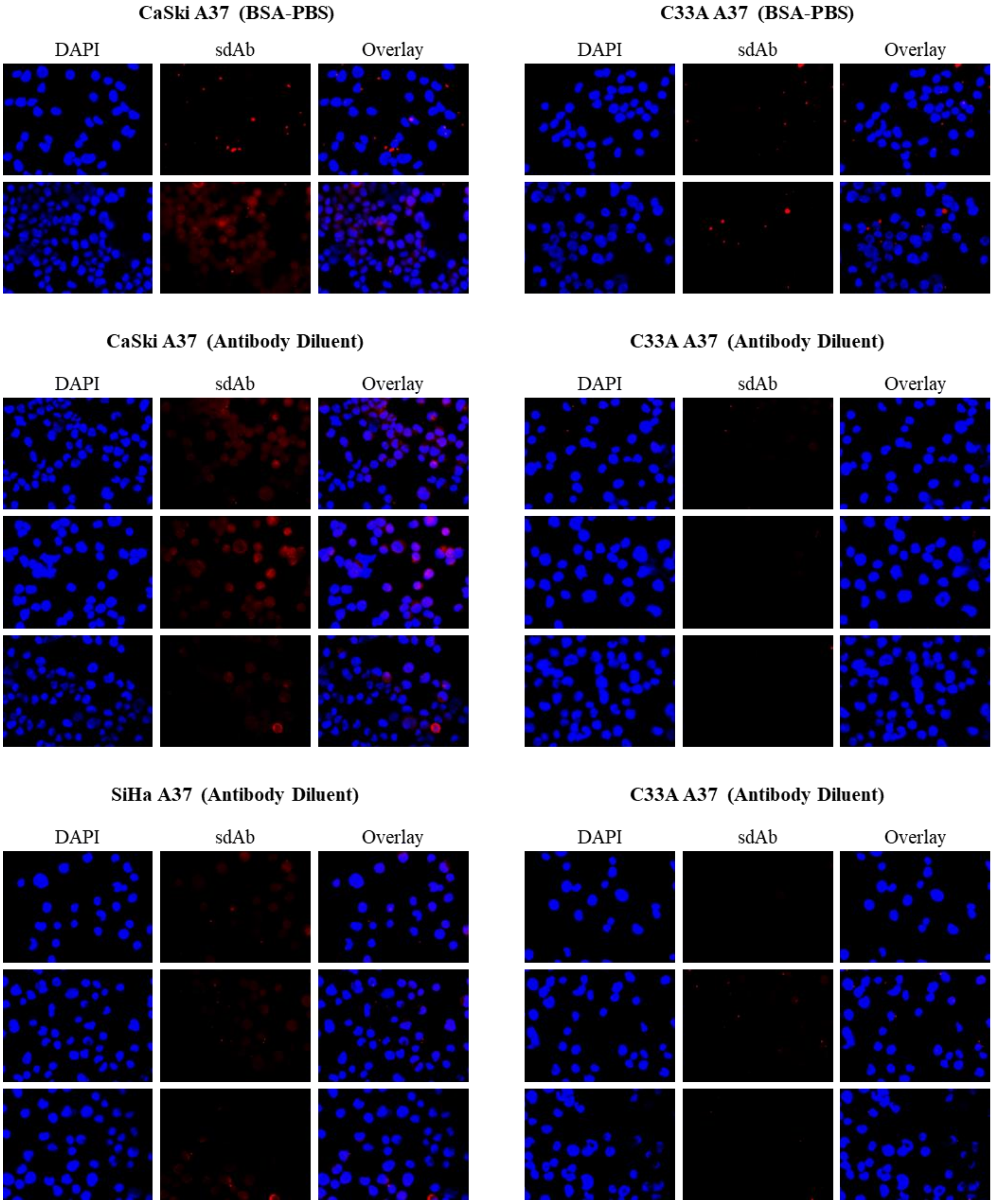

### Supplemental IF: 2A04

**CaSki 2A04 (BSA-PBS)**

DAPI sdAb Overlay

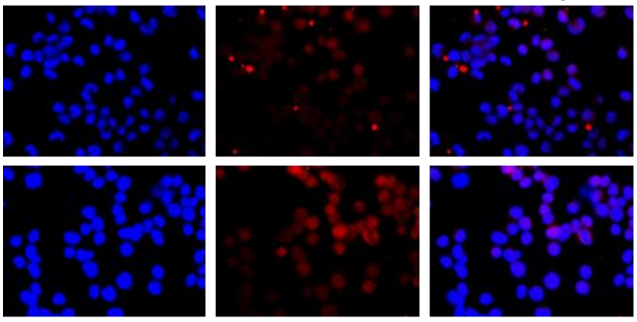

**C33A 2A04 (BSA-PBS)**

DAPI sdAb Overlay

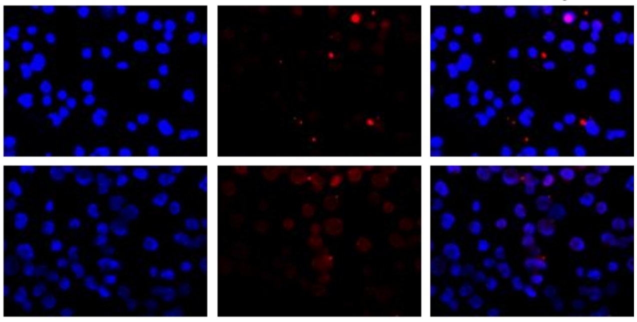

**CaSki 2A04 (Antibody Diluent)**

DAPI sdAb Overlay

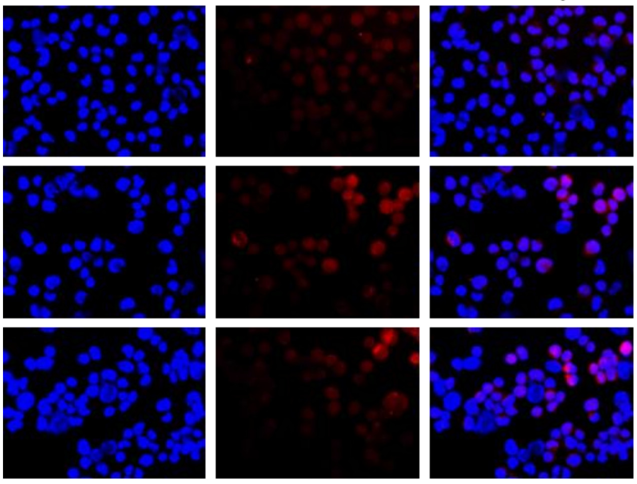

**C33A 2A04 (Antibody Diluent)**

DAPI sdAb Overlay

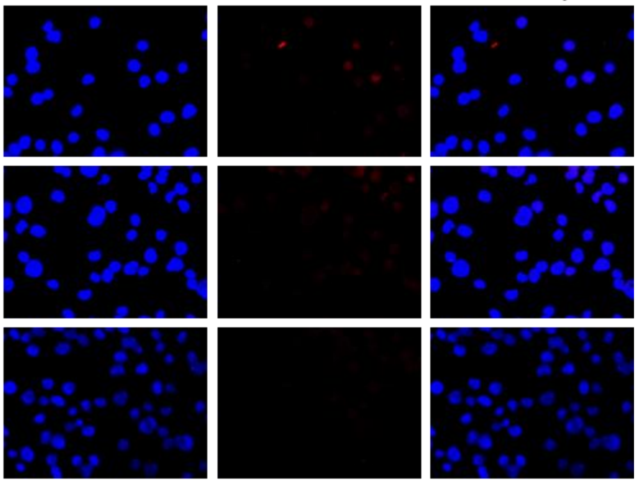

**SiHa 2A04 (Antibody Diluent)**

DAPI sdAb Overlay

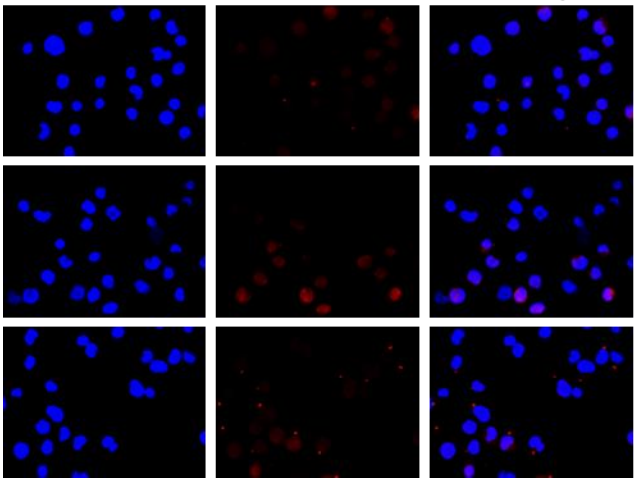

**C33A 2A04 (Antibody Diluent)**

DAPI sdAb Overlay

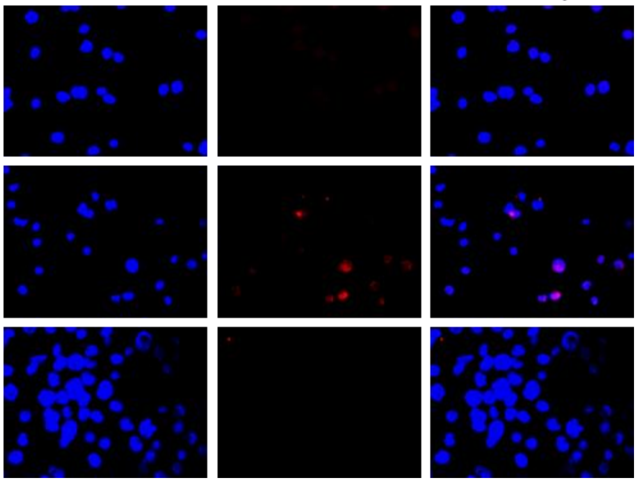

### Supplemental IF: 2A03

**CaSki 2A03 (BSA-PBS)**

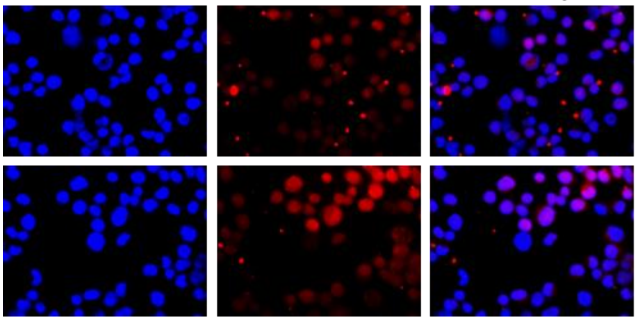


**C33A 2A03 (BSA-PBS)**

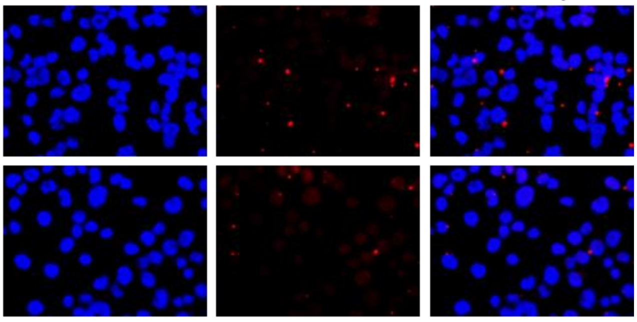


**CaSki 2A03 (Antibody Diluent)**

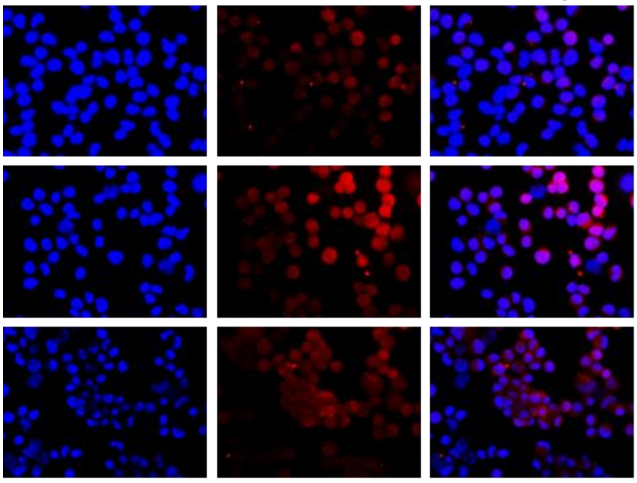


**C33A 2A03 (Antibody Diluent)**

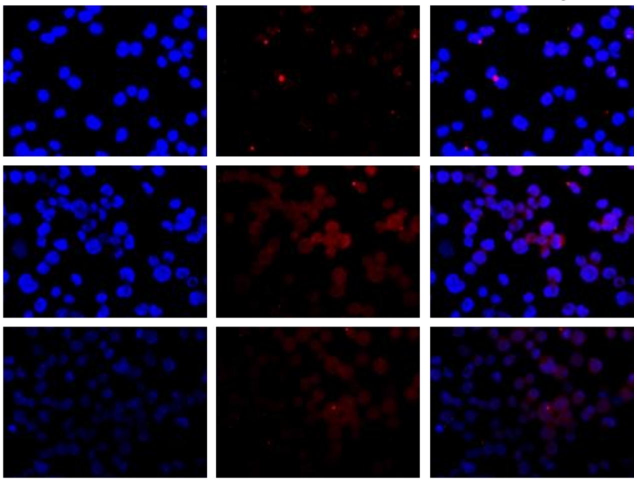


**SiHa 2A03 (Antibody Diluent)**

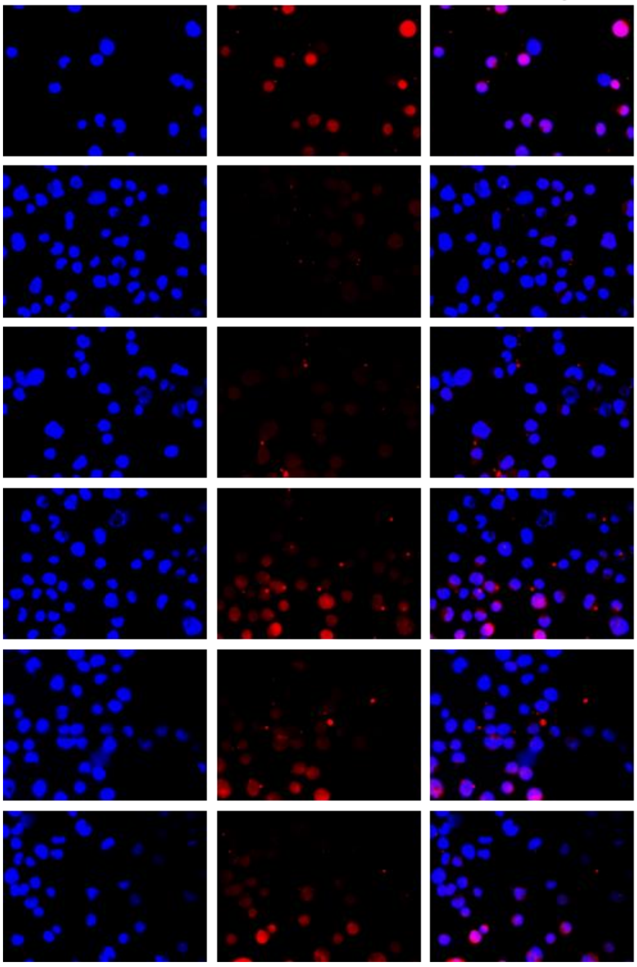


**C33A 2A03 (Antibody Diluent)**

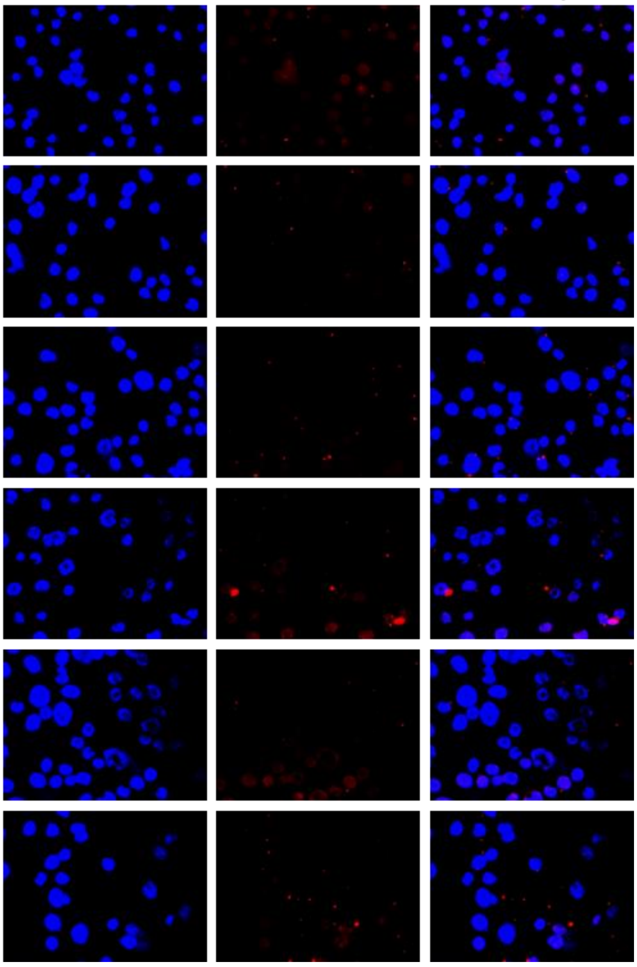

### Supplemental IF: A46*

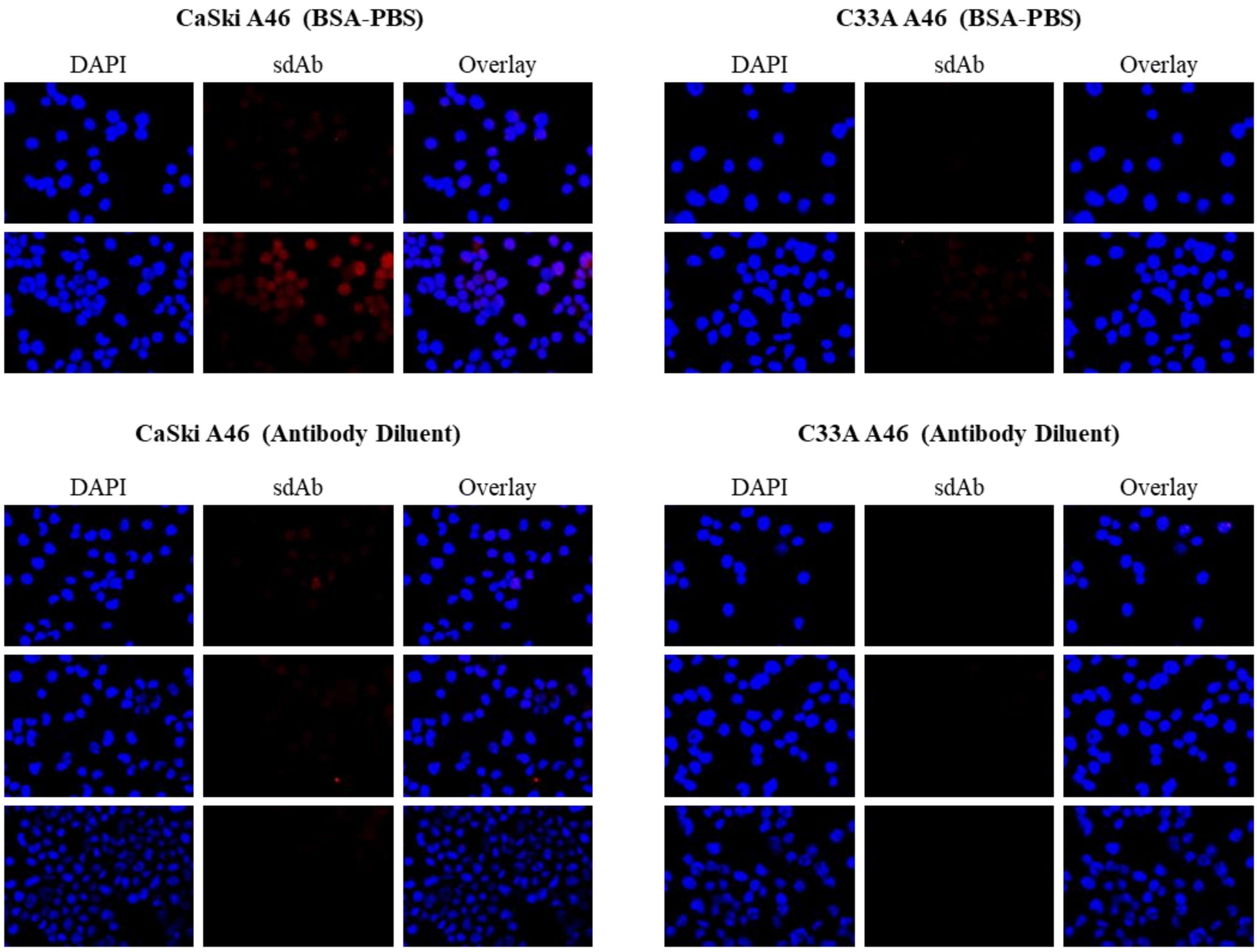


* Due to inconsistencies in results obtained with A46, the sdAb was not further investigated with SiHa cells

### Supplemental IF: A05**

**CaSki A05 (BSA-PBS)**

DAPI | sdAb | Overlay

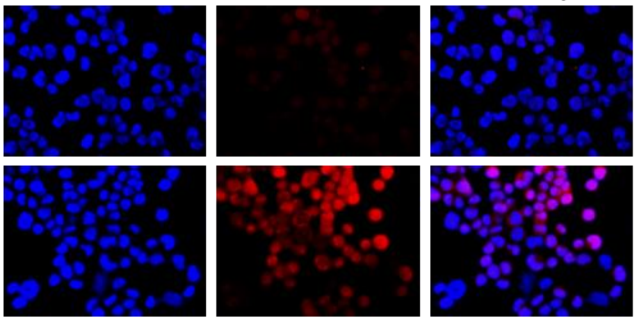

**C33A A05 (BSA-PBS)**

DAPI | sdAb | Overlay

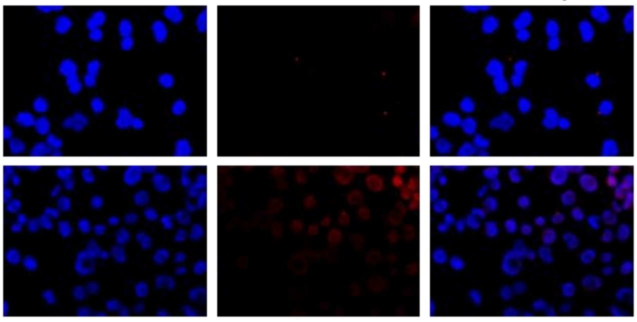

**CaSki A05 (Antibody Diluent)**

DAPI | sdAb | Overlay

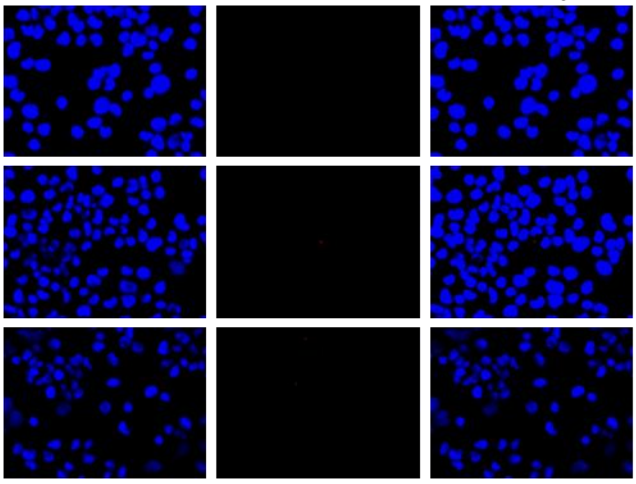

**C33A A05 (Antibody Diluent)**

DAPI | sdAb | Overlay

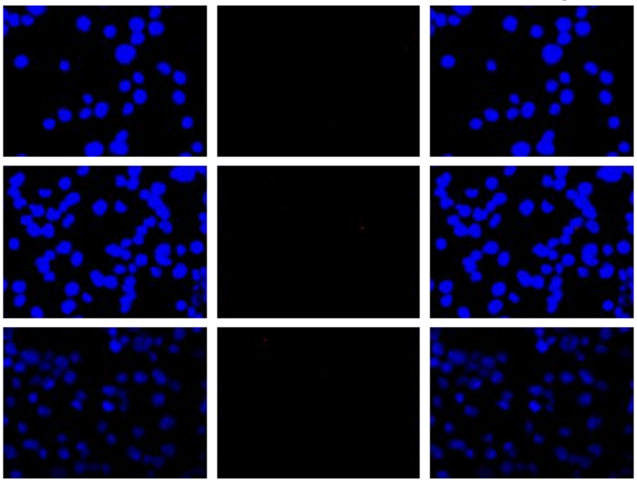

** Due to inconsistencies in results obtained with A05, the sdAb was not further investigated with SiHa cells

### Supplemental IF: 2A10***

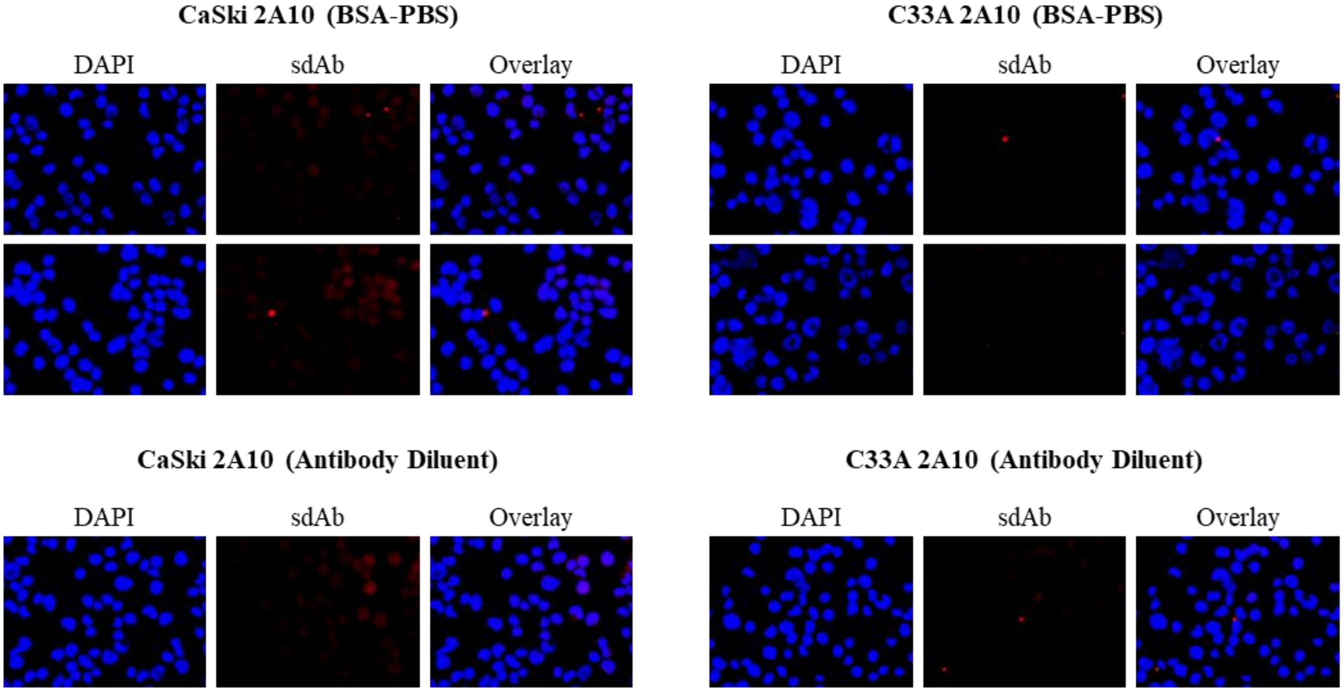


*** The antibody was not further investigated due to the lack of signal in CaSki using 2A10 and the limited amount of soluble 2A10 sdAbs.

### Supplemental IF: 2A15****

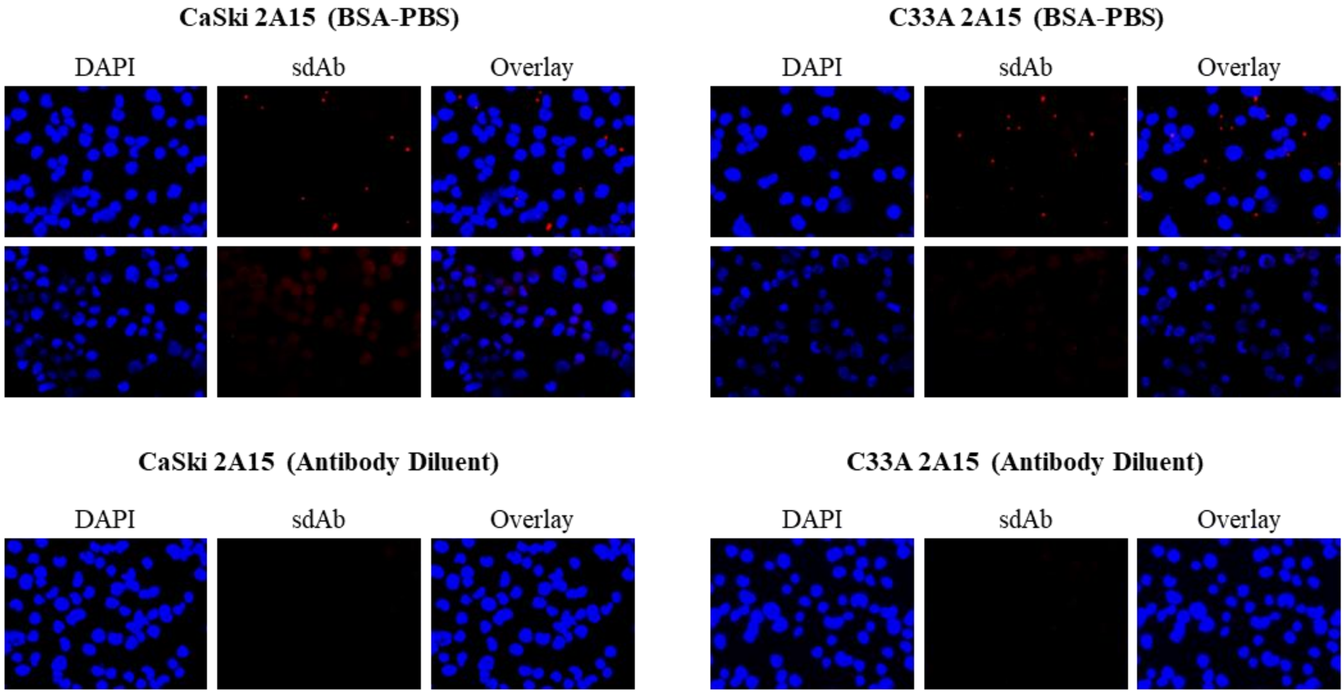


**** The antibody was not further investigated due to the lack of signal in CaSki in the initial experiments.

#### Supplemental IF: A47*****

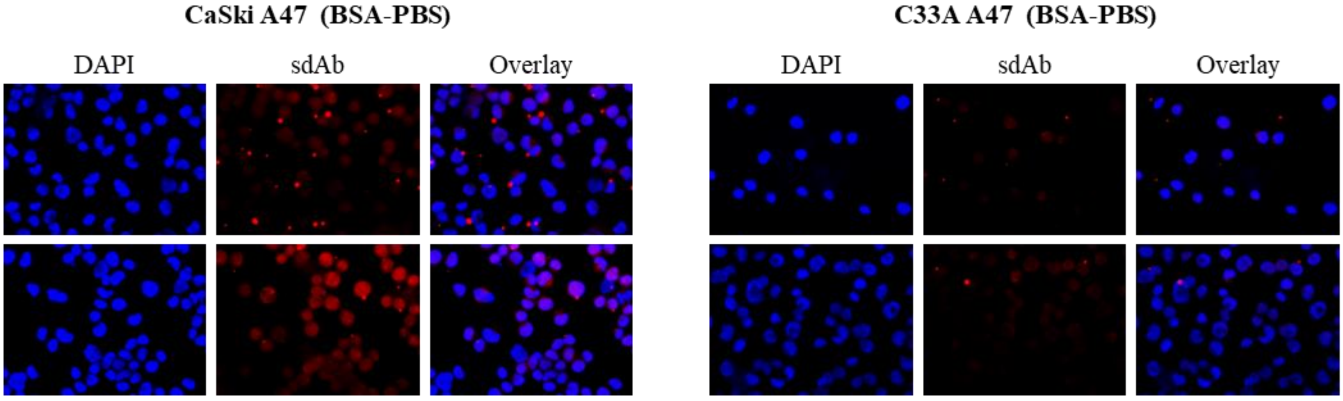


***** The antibody was not further investigated using DAKO antibody diluent as no more soluble A47 was available to complete the experiments.
