## Supplemental IF p53 and PARP for "Targeting the Human Papillomavirus 16 E6 Oncoprotein with Antibodies"

### *Evaluation of p53 restoration and PARP-1 activation*

Chamber slides (Lab-Tek, Cat# 177402) were pre-treated with Poly L Lysine before CaSki, C33A (24 000), and SiHa (12 000) cells were seeded and allowed to grow for 48-hours. Cells were transfected with 25 ng of C26, A37, C26 + A37, or mCherry mRNA using the Mirus *Trans*-IT-mRNA Transfection Kit, following the supplier recommendations for a 48-well plate (same surface area as the chamber slides). After the 4-hour transfection, the medium was replaced as before, and the cells were allowed to grow for 48-hours to reach 80% confluence. The slides were prepared as described in *Microscopy imaging* with the following modifications. Blocking and antibody incubation were performed in 4% BSA-PBS solution. P53 and PARP-1 were detected using the following antibodies: p53 MAb [DO1] (1:400, Invitrogen by Thermo Fisher Scientific, Stock Concentration: 0.2 mg/mL, Cat # AHO0152), mouse mAb to cleaved PARP [4B5BD2] (1:760, Abcam, Stock Concentration: 0.76 mg/mL, Cat # ab110315). Co-staining of the sdAbs and p53 or the sdAbs and PARP-1 was performed using the following antibodies: 1/400 dilution of Alexa Fluor 594 AffinPure Goat anti-Alpaca VHH (Jackson Immunoresearch Inc., Cat #: 128- 585-230), 1:400 Alexa Fluor 488 Donkey anti-mouse IgG (Invitrogen by Thermo Fisher Scientific, Cat# A21202). Cells transfected with the mRNA storage buffer and un-transfected cells were used as controls as before. Additionally, cells treated with Actinomycin D (Sigma, Cat# A9415) (CaSki 500 nM, SiHa and C33A 50 nM) were used as a positive control for p53 restoration and PARP-1 activation. CellProfiler (Version 4.2.1) was used to measure the amount of sdAb, mCherry, p53, and PARP-1 in the nucleus of transfected cells. The percentage of sdAb, mCherry, p53, and PARP-1 positive cells as well as the overall signal intensity was similarly evaluated as previously described (Togtema et al. 2012).

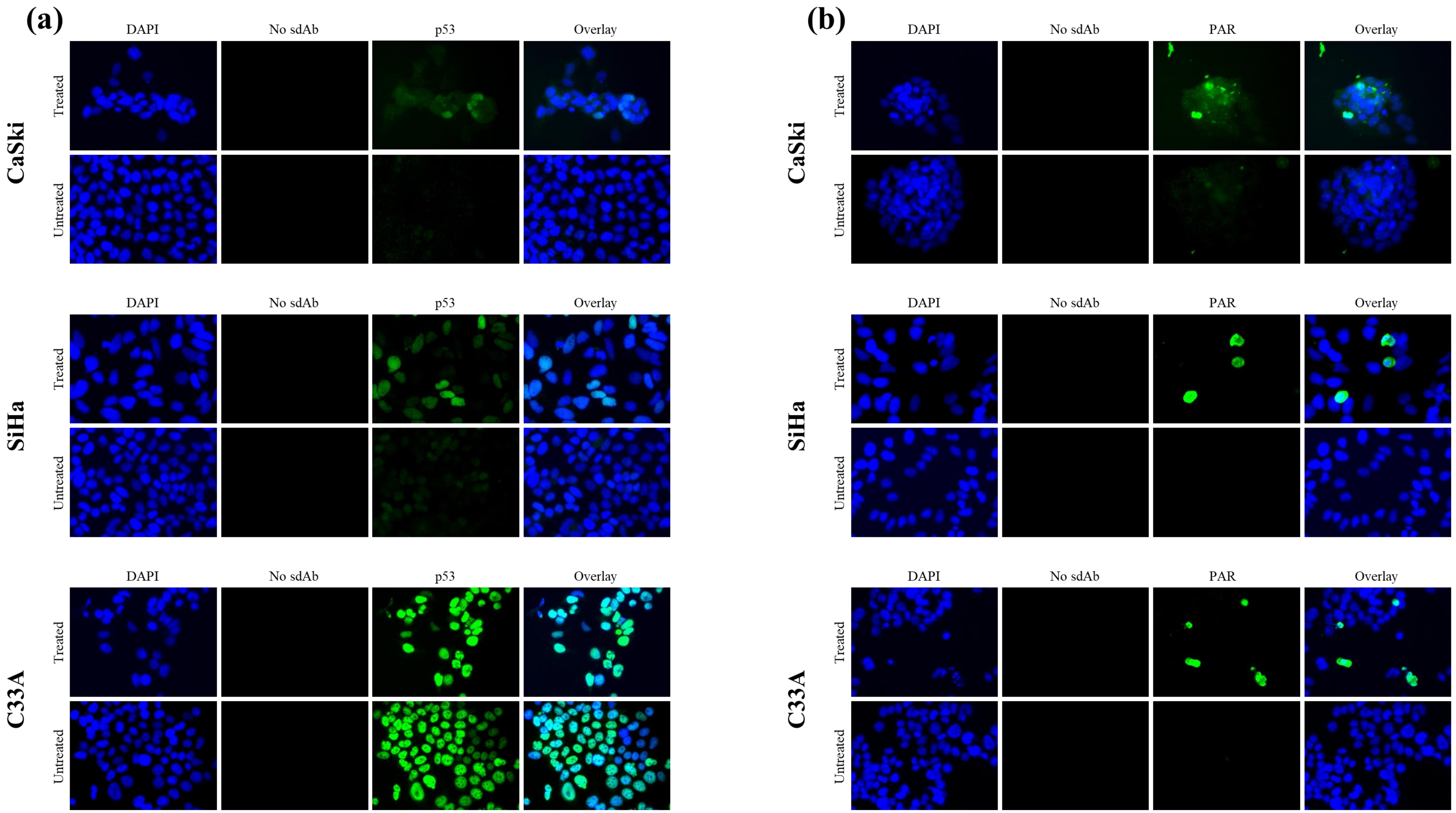


**Immunodetection of p53 and PARP in CaSki, SiHa, and C33A cells 24-hours after treatment with Actinomycin D**. 12 000 – 24 000 cells were seeded 24-hours prior to treatment. After 24-hours, existing media was replaced with 300 µL of Actinomycin D treatment solution (0.5 µL/mL for CaSki and 0.05 µL/mL for SiHa and C33A) and incubated for 24-hours. Following treatment, slides were prepared for analysis with p53 probed using 1:400 DO1 p53 monoclonal antibody, PARP probed using 1:760 Mouse mAb to cleaved PARP [4B5BD2], and both were detected using 1:400 Alexa Fluor 488 Donkey anti-mouse IgG. Nuclei were visualized using DAPI followed by imaging with the Zeiss Axiovert 200 fluorescence microscope equipped with an LD A-Plan 40x/0.50 Ph2 objective and a CCD camera with 12-bit capability and edited using ImageJ software version1.53a.

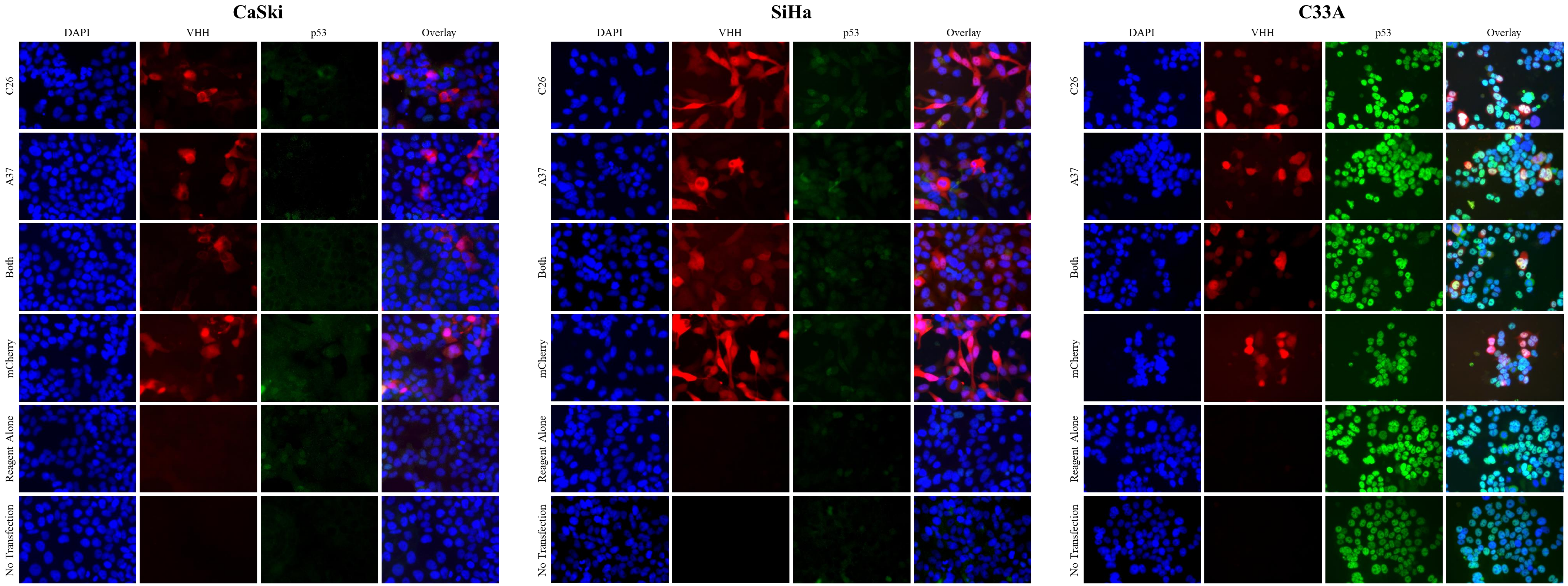


**Immunodetection of 1X (250 ng) C26, A37, and mCherry (red) and related p53 expression (green) in CaSki, SiHa, C33A cell lines 48-hours after mRNA transfection.** 12 000 – 24 000 cells were seeded 48-hours prior to transfection, with fresh media added 4-hours post transfection. Transfection was performed using the Mirus *Trans*IT-mRNA Transfection Kit following manufacturer protocols. 48-hours post transfection, slides were prepared for analysis. sdAbs were detected using 1:400 Alexa Fluor 594 + while p53 was probed using 1:400 DO1 p53 monoclonal antibody and detected using 1:400 Alexa Fluor 488 Donkey anti-mouse IgG. Nuclei were visualized using DAPI followed by imaging with the Zeiss Axiovert 200 fluorescence microscope equipped with an LD A-Plan 40x/0.50 Ph2 objective and a CCD camera with 12-bit capability and edited using ImageJ software version1.53a

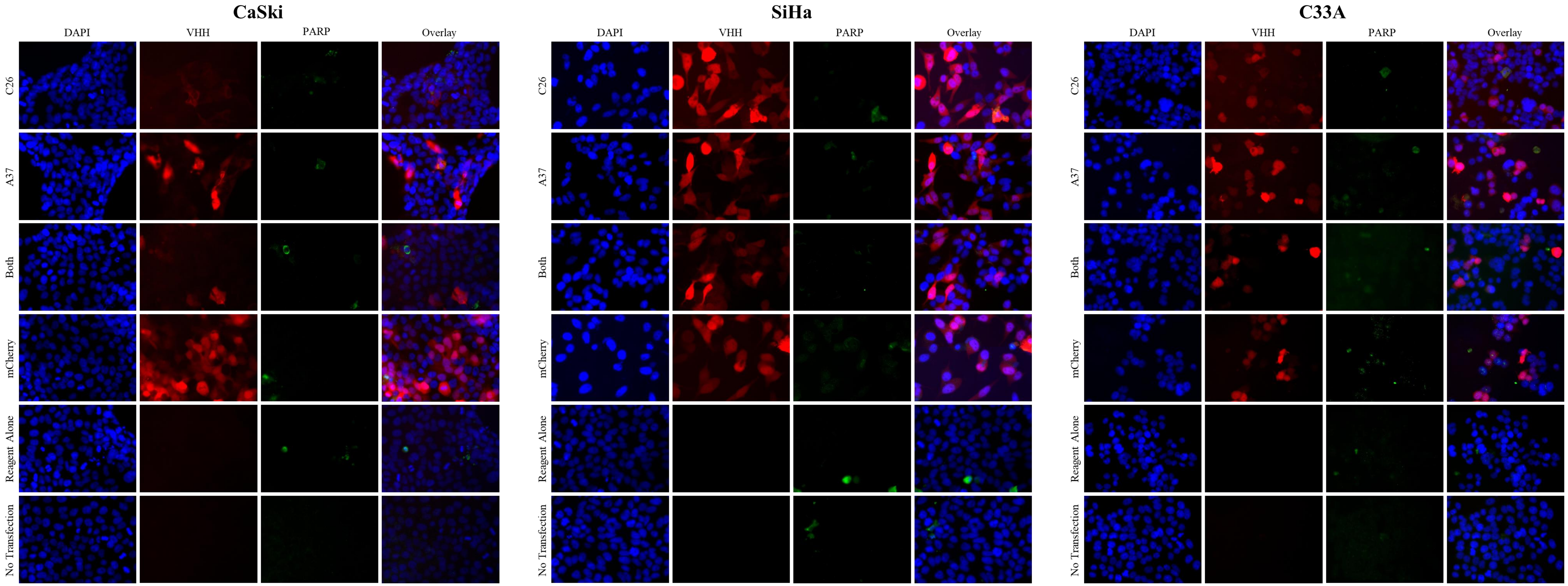


**Immunodetection of 1X (250 ng) C26, A37, and mCherry and related PARP expression in CaSki, SiHa, C33A cell lines 48-hours after mRNA transfection.**
12 000 – 24 000 cells were seeded 48-hours prior to transfection, with fresh media added 4-hours post transfection. Transfection was performed using the Mirus *Trans*IT-mRNA Transfection Kit following manufacturer protocols. 48-hours post transfection, slides were prepared for analysis. sdAbs were detected using 1:400 Alexa Fluor 594 + while PARP was probed using 1:760 Ms mAb to cleaved PARP [4B5BD2] and detected using 1:400 Alexa Fluor 488 Donkey anti-mouse IgG. Nuclei were visualized using DAPI followed by imaging with the Zeiss Axiovert 200 fluorescence microscope equipped with an LD A-Plan 40x/0.50 Ph2 objective and a CCD camera with 12-bit capability and edited using ImageJ software version1.53a.
