## Supplementary material for "Targeting the Human Papillomavirus 16 E6 Oncoprotein with Antibodies": sdAb Protein Sequences

>A01
QVQLVESGGGLVQPGGSLRLSCAASGGSFSSYALAWFRQAPGKEREFVAAVTWNGASTYYADSVKGRFTI
SRDNAKNTVYLQMNSLKPEDTAVYYCAAGPRGRYFYTAHREYAYWGQGTQVTVSSGSYPYDVPDYAGSHH
HHHH

>A05
QVQLVESGGGLVQPGGSLRLSCAVSGTTLDEYAIGWFRQAPGKEREGISCISTNGPTYYTDSVKGRFTIS
SDNAQNTVYLQMNSLKSEDTAVYYCATVGPVAWGACLPLASDDNYWGQGTQVTVSSGSYPYDVPDYAGSH
HHHHH

>A09
QVQLVESGGGLVQPGGSLKLSCAASGSISRINVMGWYRQTPGKQRELVAEITSGGSTNYVDSVKGRFTIS
RDNAKNTVYLQMNSLKPEDTAVYYCTADRFGGGSYPQREDGYDYWGQGTQVTVSSGSYPYDVPDYAGSHH
HHHH

>A27
QVQLVESGGGLVQSGGSLRLSCAASGFSFDDYAIGWFRQAPGKEREGVACILSSDGSTYYANSVKGRFTI
SSDNAKNTVYLQIDSLKPEDTAVYYCAADPHPLCGSSLVRFRGSQYDYWGQGTQVTVSSGSYPYDVPDYA
GSHHHHHH

>A34
QVQLVESGGGLVQAGGSLKLSCAAVGGSVFSRPTMGWYRQIPGKQREQRDLVAIITSGGDTTYADSVKGR
FSISRDKAKKTYLQMNNLKPEDTGIYYCNARSSTYSSSNIWGQGTQVTVSSGSYPYDVPDYAGSHHHHHH

>A37
QVKLEESGGGLVQAGGSLRLSCVASGSTFIITDMAWYRQAPGKQRELVAGITLRGGTNYADSVKGRFTIS
RDNVKNTVTLQMNSLKPEDTAVYYCNAKVAEWRNRPDYWGQGTQVTVSSGSYOYDVODYAGSHHHHHH

>A45
QVQLVESGGGLVQAGGSLRLSCAASGFSFDDYAIGWFRQAPGKEREGVACTNSKYGDTYYAEAVKGRFTI
SSDNAKNTVYLQMNSLKPEDTAVYYCVADQSLGCGVGDFWGQGTQVTVSSGSYPYDVPDYAGSHHHHHH

>A46
QVKLEESGGGLVQAGGSLRLSCAASGSILSIDDMGWYRQAPGKQRELVASITSDGSTNYADSVKGRFTIS
RDNAKNTVYLQMNRLKPEDTAVYYCNADLIPYSDYALPSYWGQGTQVTVSSGSYPYDVPDYAGSHHHHHH

>A47
QVQLVESGGGLVQAGGSLRLLCVVSGSIFNIKTVGWYRQAPGKERELVADIRTSGTQTQYADFAKGRFTI
SRDNGGRKVYLEMSDLKPEDTAVYYCRAERWTLSAGTGDYWGQGTQTVSSGSYPYDVPDYAGSHHHHHH

>C11
QVQLVESGGGLVQAGGSLRLLCVVSGSISNIKTVGWYRQAPGKQREFVADIVISGSKTQYADSVKGRFTI
SRDNSGRKVSLEMSDLKPEDTAVYVCRAERWSLSAGTGDYWGQGTQVTVSSGSYPYDPDYAGSHHHHHH

>C36
QVQLVESGGGLVQPGGSLRLSCAASGFTFDGFVMSWVRAQPGKQREYVAAILRNGRTNYADSVKGRFTIS
RDNAKNTMYLQMNNLKPEDTAVYYCGAAIPRRPGEDLNGYDYWGQGTQVTVSSGSYPYDVPDYAGSHHHH
HH

>C38
QVQLVESGGGVVQAGGSLRLSCAPSGNIFSINTMGWHRQAPGKQREFVARIRSSGQTNYADSVKGRFTIS
KDSAKDTVYLQMDNLQPEDSAVYYCTFNSGGWGVERKRRDYWGQGTQVTVSSGSYPYDVPDYAGSHHHHH
H

>2A03
QVQLVESGGGLVQAGGSLRLSCAASGRTSSINIMGWYRQPPGKQREMVATIATGGTTNYAESVKGRFTIS
RDGAKFVYLQLNDLKPEDTAVYYCNAYRYAVGKNQRAWDIWGQGTQVTVSSGSYPYDVPDYAGSHHHHHH

>2A04
QVKLEESGGGVVQAGGSLRLSCAPSGNIFSINTMGWHRQAPGKQREFVARIRSSGQTNYADSVKGRFTIS
KDSAKDTVYLQMDNLQPEDTAVYYCTFNTGGWGVERQRRDYWGQGTQVTVSSGSYPYDVPDYAGSHHHHH
H

>2A10
QVQLVESGGGLVQAGGSLKLSCAAVGGSVFSRPTMGWYRQIPGKQRELVATITNDGSTYYEEAVKGRFTI
SRDNAKNTLSLQMNSLKPEDTAVYYCSTRSSWGQGTQVTVSSGSYPYDVPDYAGSHHHHHH

>2A12
QVKLEESGGGLVQPGGSLRLSCAGSGTIVYIHAMGWYRRAPGSERLVATIARDGTTHYADSVKGRFTISR
DNDRNMMWLQMNSLKPEDTAVYYCNADLVERSWGIRRDYWGQGTQVTVSSGSYPYDVPDYAGSHHHHHH

>2A15
QVQLVESGGGVVQAGGSLRLSCAPSGNIFSINTMGWHRQAPGKQREFVARIRSSGQTNYADSVKGRFTIS
KDSAKDTVYLQMDNLQPEDSAVYYCTFNSGGWGVERERRDYWGQGTQVTVSSGSYPYDVPDYAGSHHHHH
H

>2A17
QVQLVESGGGLVQAGGSLRLSCEASGNIRSLGVMGWYRQAPGKQRELVADITVWRTTNYGDSVKGRFTIS
RDNANTIFLQMNSLKPEDTSLYYCNYRDSRDYWGQGTQVTVSSGSYPYDVPDYAGSHHHHHH

>2A51
QVQLVESGGGLVQDGGSLRLSCEASGLPFRDNAMNWYRQAPTGKQRDFVARISRGGSTKYAEFVKGRFAI
SRDNAKNTVALQMNSLKPEDTAVYYCFAEGPPGNVWGQGTQVTVSSGSYPYDVPDYAGSHHHHHH

>2A78
QVQLVESGGGVVQAGGSLRLSCAPSGNIFAINSMGWHRQAPGKQRELVATIRSSGKTNYADSVKGRFTIS
KDSSKDTVFLQMDNVQPEDTAVYYCTFNTGGWGVQRTRRDYWGQGTQVTVSSGSYPYDVPDYAGSHHHHH
H

>C26
QVQLVESGGGLVQAGGSLRLSCAASGSIYSINAMGWYRQAPGKQRELVAVITSSGSTNYADAVKGRFTIS
RDNAKNQVYLQMNSLKPEDTAVYYCHAWSPFRGSFSGDALDAWGQGTQVTVSSGSYPYDVPDYAGSHHHH
HH
